# Intraspecific Variation in Microsatellite Mutation Profiles in Daphnia magna

**DOI:** 10.1101/540773

**Authors:** Eddie K. H. Ho, Fenner Macrae, Leigh C. Latta, Maia J. Benner, Cheng Sun, Dieter Ebert, Sarah Schaack

## Abstract

Microsatellite loci (tandem repeats of short nucleotide motifs) are highly abundant in eukaryotic genomes and are often used as genetic markers because they can exhibit variation both within and between populations. Although widely recognized for their mutability and utility, the mutation rates of microsatellites have only been empirically estimated in a few species and have rarely been compared across genotypes and populations and intraspecific differences in overall microsatellite content have rarely been explored. To investigate the accumulation of microsatellite DNA over long-and short-time periods, we quantified the abundance and genome-wide mutation rates in whole-genome sequences of 47 mutation accumulation (MA) lines and 12 non-MA lines derived from six different genotypes of the crustacean *Daphnia magna* collected from three populations (Finland, Germany, and Israel). Each genotype possessed a distinctive microsatellite profile and clustered according to their population of origin. During the period of mutation accumulation, we observed very high microsatellite mutation rates (a net change of −0.19 to 0.33 per copy per generation), which surpass rates reported from a closely-related congener, *D. pulex,* by an order of magnitude. Rates vary between microsatellite motifs and among genotypes, with those starting with high microsatellite content exhibiting greater losses and those with low microsatellite content exhibiting greater gains. Our results show that microsatellite mutation rates depend both on characteristics of the microsatellites and the genomic background. These context-dependent mutation dynamics may, in conjunction with other evolutionary forces that may differ among populations, explain the differential accumulation of repeat content in the genome over long time periods.

## Introduction

Microsatellite loci, also known as short tandem repeats, are repetitive regions of the genome known for their propensity to mutate rapidly (e.g., Sun et al., 2012). Although exact definitions of microsatellites vary, they typically involve tandem arrays of short motifs (typically, 1-6 bp long, although longer motifs can also be found in tandem arrays). Microsatellites can be located inside or outside coding regions of the genome, and have been shown to influence a range of phenotypes from gene expression to genetic disease (Feupe Fotsing et al., 2018). Previous reports of microsatellite mutation rates (MMRs) have consistently shown them to be higher than substitution rates in unique sequence, often by several orders of magnitude (reviewed in Ellegren, 2004). Because of their mutability, microsatellites have frequently been used in population genetics studies and there is increasing interest in the role they may play in adaptation, plasticity, and disease (Haasl and Payseur, 2013; Hannan, 2018).

There are three mechanisms of mutation that have been proposed to explain the patterns of higher mutation rates at microsatellite loci: retrotransposition, unequal crossing over, and DNA slippage. Retrotransposition, in particular, could explain the frequent observation that microsatellites tend to be A-rich, although it is less clear how retrotransposition would impact mutation rates of microsatellites once they are formed. Unequal crossing over is thought to increase in frequency at repeat-rich loci and can, potentially, lead to the expansion or contraction of tandem arrays with equal probability. The most often discussed mechanisms of microsatellite mutation is strand slippage during DNA replication and repair (Kornberg et al., 1964), whereby the array of repeats can cause potential mispairing between template and nascent strands of DNA. If uncorrected by DNA repair mechanisms, slippage can lead to the expansion or contraction of a tandem array and may do so in a motif-or length-dependent manner (Eckert and Hile, 2009). When substitutions occur at microsatellite repeats, they result in the loss (or ‘death’) of the repeat, in addition to loss or contraction due to deletions or contractions during slippage (Kelkar et al., 2011). A given microsatellite locus can experience any of a number of different types of mutation (e.g., insertions, deletions, duplications, slippage, and substitutions) which can result in either an expansion or contraction of that tandem array, the interruption of the array, or the increase or decrease in copy number of the array. Because all these mutation types will contribute to overall copy number for any given motif (referred to as a kmer, hereafter), genome-wide analyses of microsatellite mutation rates can benefit from looking at rates of copy number increase and decrease as a global metric of the impact of mutation at these loci.

Microsatellite mutation rate (MMR) variation based on the composition of the motif (AT/GC content), length of the motif (unit length; e.g., dinucleotide versus trinucleotide repeats), and the length of the array (e.g., the number of repeats occurring in tandem at a given locus) has been the focus of previous studies in a variety of systems (reviewed in Bhargava and Fuentes, 2010). Theoretically, mutation rates would be expected (A) to be higher in AT-rich regions (due to the lower number of hydrogen bonds between base pairs), (B) to decrease as a function of unit length based on the strand slippage model of mutation, and (C) to increase as a function of array length, given the increased number of targets for mutation (reviewed in Eckert and Hile 2009). Indeed, empirical studies have shown that microsatellites with high AT-content tend to mutate at higher rates than those that are GC-rich and that di-nucleotide rates are higher than tri-nucleotide repeats (Chakraborty et al., 1997). Rates of expansion versus contraction, however, have been shown to depend on starting length, with shorter arrays tending to increase in length and longer arrays tending to decrease in length (Lai and Sun, 2003; Seyfert et al., 2008). Indeed, if MMRs vary based on any of these factors, one could make predictions about the accumulation of microsatellites across the genome over long time periods based on starting composition of the repeat content.

As with most types of mutations, mutation rate estimates are typically performed on one or a few genotypes for a representative model species, and then used to extrapolate mutation rate estimates for congeners, or even more widely, despite a lack of evidence for generalizing to this degree (e.g., mutation rate estimates for *Drosophila melanogaster* are routinely used as a proxy for all insects, despite known variation in rates estimates between genotypes (Haag-Liautard et al., 2007)). The degree to which microsatellite mutation rates and patterns of microsatellite accumulation vary between genotypes and populations, intraspecifically, or among closely-related species with similar lifespans, physiologies, and life histories has remained largely unexplored. Given that the rate of mutation itself is a trait that can evolve, knowing the level of intraspecific variation upon which evolutionary forces can act to increase or decrease the rate over time (Lynch, 2010), as well as knowing what factors influence rate differences, is of major interest to biologists (Baer et al., 2007). Most recently, it has been proposed that mutation rates across species hover near a “drift barrier”, meaning that they are only driven down by selection to the extent possible based on the effective population size, at which point they can not be lowered further due to the relative power of genetic drift which permits mutations that increase (or maintain) the rate (Lynch, 2010). Knowing the level of intraspecific variation in mutation rates is essential for assessing the potential of a drift barrier to explain mutation rate variation within and between species. Mutation accumulation (MA) experiments provide the least biased estimates of mutation rates available (Halligan and Keightley, 2008), although they can only be conducted in organisms that can be reared in a controlled environment with short generation times.

Here, we present data from 6 genotypes of *Daphnia magna*—2 each from three populations (Finland, Germany and Israel), and compare our results to previously published estimates of MMR in the congener, *D. pulex* (Flynn et al., 2017). *D. magna* is an important model organism for ecology, evolutionary biology, and genomics studies (Miner Brooks E. et al., 2012; Schaack, 2008). The cyclical parthenogenetic nature of *Daphnia* makes them an ideal organism to use in MA experiments because clonal reproduction facilitates their long-term maintenance in the lab. Our goal is to characterize both the microsatellite landscape and mutational profiles across these 6 genotypes in order to determine if there is a relationship between the two, and to assess the degree to which they may vary among genotypes, populations, and closely-related species. In addition, we report the microsatellites that are most abundant and most mutable to determine if there are features of individual microsatellites (unit length or content) that determine differences in mutation dynamics among loci. Identifying patterns using mutation accumulation data collected on experimental time-scales where selection is minimized and contrasting such data with patterns of microsatellite accumulation over long time-periods can reveal the degree to which evolutionary forces are shaping microsatellite landscapes in nature.

## Methods

### Study System

The *D. magna* genotypes used in this experiment were collected along a latitudinal gradient that captures a range of environmental variation including temperature and photoperiod. Specifically, two unique genotypes from each of three populations (Finland, Germany, and Israel) were used to initiate the control and mutant lines. The stock cultures for each genotype were maintained in 250 mL beakers containing 175-200 mL of Aachener Daphnien Medium (ADaM; Klüttgen et al., 1994) under a constant photoperiod (16L:8D) and temperature (18 °C), and fed the unicellular green alga *Scenedesmus obliquus* ad libitum (2-3 times per week).

Two types of controls were used in this experiment. First, we established and maintained populations of large size from descendants of the same genotypes used to initiate the MA lines. In these large population controls, mutations may occur, but are more likely to be purged by selection due to competition among clones (relative to the MA lines, where clones are reared individually and experience no competition). At the end of the mutation accumulation period, these large populations are sampled and DNA is extracted (referred to hereafter as ‘extant controls’ or ECs). Although mutations can occur in these lineages, the paucity of mutations observed is consistent with the idea that the MA protocol (bottlenecking lineages each generation by transferring a single individual) does, indeed, minimize selection (see below). The second set of control lines sequenced was from tissue harvested from immediate descendants of the individual from which progeny were used to establish the MA lines (referred to hereafter as ‘starting controls’ or SCs) at the beginning of the mutation accumulation period.

### Mutation Accumulation Experiment

Starting control (SC), extant control (EC), and mutation accumulation (MA) lines were initiated from clonally-produced offspring of a single asexual female isolated from the stock cultures of each of the 6 genotypes described above. Tissue samples for SCs consisted of 5-20 individual *Daphnia* collected from each genotype beginning two weeks after initiation of the MA experiment. Individuals were placed in a 1.5 mL microcentrifuge tube, frozen in liquid nitrogen, and stored at −80 °C until DNA extraction. Two EC lines were initiated from each SC, and were sampled at the end of the experiment (approximately 2.5 years after initiation of the experiment). Two EC lines were maintained in separate 3 L jars containing 2 L of ADaM, under constant temperature (18 °C) and photoperiod (16L:8D), and fed the unicellular green alga *S. obliquus* ad libitum. The media in the jars was replaced every 2-3 weeks, and individuals in the two replicate jars were mixed to maintain as much genetic homogeneity among the jars as possible. The maintenance of EC lines in large jars ensures that population densities, which varied between several hundred to a few thousand individuals, were high enough that new mutations with deleterious effects should be efficiently eliminated from the populations by purifying selection.

In addition to the control lines, between 10-15 MA lines were initiated from each of the 6 starting genotypes. The MA protocol used here has been described previously (Eberle et al., 2018). Briefly, MA lines were initiated by placing a single clonally-produced female in a 250 mL beaker containing 100 mL of ADaM supplemented with *S. obliquus* at a concentration of 600,000 cells/mL. All MA lines were maintained in environmental conditions identical to the control lines (16L:8D, 18 °C). The food/media mixture in each beaker was replaced once per week, and each line was fed a prescribed volume of concentrated *S. obliquus* three days after the media replacement to reset the algal cell concentration in the beaker to 600,000 cells/mL. From generation to generation, each MA line was propagated via single progeny descent by taking a single juvenile offspring from the second clutch of the previous generation. A series of backups were maintained in parallel with the focal lineages in the event that the single individual intended to be used to establish the next generation died before reproduction, or was a male. In the event that the focal lineage and all backup lineages died before reproduction or were all males, the lineage was declared extinct and a new MA line was established from the ongoing EC lines. Tissue samples for each of the MA lines were isolated every 5 generations, and at the end of the MA experiment the samples taken from lines with the greatest number of generations of mutation accumulation were used for DNA extraction and sequencing (2-26 generations, with an average of 12.4 generations per line).

### DNA Extraction and Sequencing

Five clonal individuals from each MA line and controls (1 starting control [SC] and 2 extant controls [EC] per genotype) were flash frozen for DNA extractions. DNA was extracted (2 extractions per line with 5 daphnia each) using the Zymo Quick-DNA Universal Solid Tissue Prep Kit (No. D4069) following the manufacturer’s protocol (DNA from a few samples was also extracted with the Qiagen DNeasy Blood and Tissue Kit, No. 69504). DNA quality was assessed by electrophoresis on 3% agarose gels and DNA concentration was determined by dsDNA HS Qubit Assay (Molecular Probes by Life Technologies, No. Q32851). The Center for Genome Research and Biocomputing at Oregon State University generated 94 Wafergen DNA 150bp paired-end libraries using the Biosystems Apollo 324 NGS library prep system. Quality was assessed using a Bioanalyzer 2100 (Agilent Technologies, No. G2939BA) and libraries were pooled based on qPCR concentrations across 16 lanes (2 runs). Libraries were sequenced on an Illumina Hiseq 3000 (150 bp PE reads) with an average insert size of ∼380bp to generate approximately 50x coverage genome-wide for each sample (Table S7).

### Tandem repeat quantification

Sequenced reads from all lines were trimmed of adapters and decontaminated to remove mitochondrial sequences. Overlapping reads were merged with BBmerge (Bushnell et al., 2017). To quantify tandem repeats, the reads were input into the program k-seek (Wei et al., 2014). The program k-seek detects tandem repeats (kmers) of 1-20 bp, requiring that the kmers repeat tandemly over at least 50 bp on a given read, allowing for one base pair mismatch per repeat unit. Offsets and reverse complements of each kmer are combined and the output is the total count across all reads for each kmer. Because k-seek has the requirement that tandem repeats span at least 50 bp, the threshold number of repeat units required for detection decreases as the length of the repeat unit increases (e.g., 1-mers require at least 50 repeats to be detected while 5-mers only require a minimum of 10 repeats to be detected). On the other hand, the maximum number of repeat units on a single read decreases as the kmer length increases. We do not observe a detection bias towards kmers with longer or shorter lengths (Table 2), which suggest that kmer length is not the main determinant of kmer detection.

To compare across samples, we normalized kmer counts by dividing copy number counts by the median sequence depth matched by the GC-content of the kmer. We first constructed *de novo* reference genomes for each of the six *D. magna* starting genotypes using Spades (Bankevich et al., 2012). Reads from each line were mapped to their corresponding reference genome using BWA default settings (Li and Durbin, 2009). Following Flynn et al. (2017), output BAM files were input into a custom script to calculate the coverage depth at each base and the GC-content of their nearby region (https://github.com/jmf422/Daphnia-MA-lines). We then group each base pair based on their nearby GC-content in the following bins: {0-0.3, 0.3-0.35, 0.35-0.4, 0.4-0.45, 0.45-0.5, 0.5-0.55, 0.55-0.6, 0.6-1}; we used wider bins for GC-content <= 0.3 and > 0.6 because there were many fewer sites containing very low and high GC-content. For each GC-content bin, we then determined the median base pair depth for use as the normalization factor. For each line, we normalize the total count of each kmer by dividing the total count by the normalization factor that corresponds with the GC-content of that kmer. This normalization approximates the copy number per 1x coverage of each kmer, which for simplicity we will refer to as the ‘copy number’.

Note that, after normalization, the total base pairs covered by a particular kmer in a particular line (i.e., kmer length * normalized copy number) can fall below 50 bp even though k-seek required tandem repeats span at least 50 bp. This is due to an over-correction by our normalization method (e.g., for a 1-mer that spans exactly 50 bp in the genome, there may be many reads that encompass the whole 50 bp array, but also many reads that only encompass a portion of the array). In this case, k-seek will not count the number of 1-mer repeats in reads that do not contain the full 50 bp array, because it falls below its threshold array length requirement. However, those reads are still counted towards our normalization factor. Due to this, the normalized copy number can fall below 50 for these 1-mers. This over-correction due to normalizing total copy number by the average (or median) coverage is present in all previous analyses that utilized k-seek (Flynn et al., 2017, 2018; Wei et al., 2014). Overall, this would cause an underestimation of the total kmer content in the genome.

### Mutation rate estimation

k-seeks outputs the total count of a given kmer across all locations in the genome as long as the requirements mentioned above are met. Thus, for our estimation of mutation rates, we define mutation as the change in the total copy number of a kmer, which could have occurred at one or more locations in the genome. For each genotype, we restrict our mutation rate analysis to kmers where the SC line had at least six copies and each of the MA lines had at least two copies. This allowed us to estimate mutation rates for 71 kmers, on average, for each of the six genotypes. We define the *genomic mutation rate of* kmer j in MA line i as U_i,j_ = (c_i,j_ - c_SC,j_)/g_i_, where c_i,j_ and c_SC,j_ represents the copy number of kmer j at MA line i and the SC line, respectively, and g_i_ represent the number of MA generations for MA line i. We found that the absolute value of the genomic mutation rate was strongly correlated with the abundance of the kmer in the SC lines (Figure S1). This was not surprising because highly abundant kmers likely represent a larger mutational target. To account for differences in the initial abundance of kmers, we define the *per copy mutation rate* of kmer j in MA line i as u_*i,j*_ = U_*i,j*_ / c_*SC,j*_. We define the overall genomic and per copy mutation for kmer j of a genotype as U_*j*_ and u_*j*_, respectively, which is calculated by taking the average U_*i,j*_ and u_*i,j*_ across all MA lines of the genotype. We calculated mutation rates for EC lines in the same way as we did for MA lines. To estimate the number of generations that each of the EC lines were maintained, we divided the length of the experiment (830 days) by their estimated generation time.

### Comparison to D. pulex

Throughout our study, we compare our *D. magna* microsatellite results to previously published results based on a dataset from *D. pulex* MA lines (Flynn et al., 2017). Briefly, Flynn et al. (2017) examined the microsatellite content from 28 MA lines and six non-MA lines that were all initially generated from a single ancestral genotype. Next generation sequencing was done following a Illumina Nextera library preparation (10x coverage, 100 bp PE reads). They analyzed kmers copies using k-seek and normalized copy number estimates as we described above (we used their study as a guide for our copy number normalization steps). In addition to shorter read lengths and lower coverage depths of sequencing, another difference in the study is the controls: they did not sequence their initial ancestral genotype, but used the average copy number of the six non-MA lines as a proxy for the ancestral state.

### Data Availability Statement

The authors affirm that all data necessary for confirming the conclusions of the article are present within the article, figures, and tables and that all sequence data generated will be submitted to GenBank upon acceptance of the article.

## Results

### Microsatellite copy number profiles in D. magna

We scanned for kmers of lengths up to 20 bp across the genome of individuals sequenced from 47 mutation accumulation lines (MA) derived from 6 starting genotypes (“starting controls” [SC]) and 12 lines (2 per starting genotype) maintained in large population sizes in parallel with the MA lines sampled at the end of the experimental period (“extant controls” [EC]). After normalization by depth of coverage, the total number of base pairs (per 1x coverage) composed of kmers ranged from 97 to 145 Kb across our six *D. magna* starting genotypes, which represented 0.07 to 0.1% of the 141 Mb *D. magna* reference genome (Figure S2). Across all SC, EC and MA lines, the median number of base pairs composed of kmers was 121 kb (0.085% of the genome). In contrast, the median kmer content of *D. pulex* was 1.2 MB (0.6% of the estimated 200 Mb *D. pulex* genome), which is an order of magnitude higher than in *D. magna* (Flynn et al., 2017). *The kmer content in our D. magna* lines was more similar to that in *Chlamydomonas reinhardtii*, which contains an average of 180 Kb (0.15% of the genome) (Flynn et al., 2018).

We performed a principal components analysis (PCA) using the copy number of the 100 kmers (average repeat unit length of 10.9) with non-zero copy numbers across all 65 lines in order to look for distinctive patterns of the microsatellite landscape across the 6 genotypes. On the first and second principle components axes, the lines clearly clustered based on their population of origin (Figure 1). We additionally performed a k-medoids analysis using the first 10 principle components (these 10 PCs explained 83% of the variation in copy number). We found that six clusters maximized the average silhouette across lines. Each of these six clusters contained the SC line, all of its descendent MA lines, and the EC of that genotype (Figure 1). Overall, the kmer copy number profiles distinguished lines based on their population and genotype.

**Figure 1.**
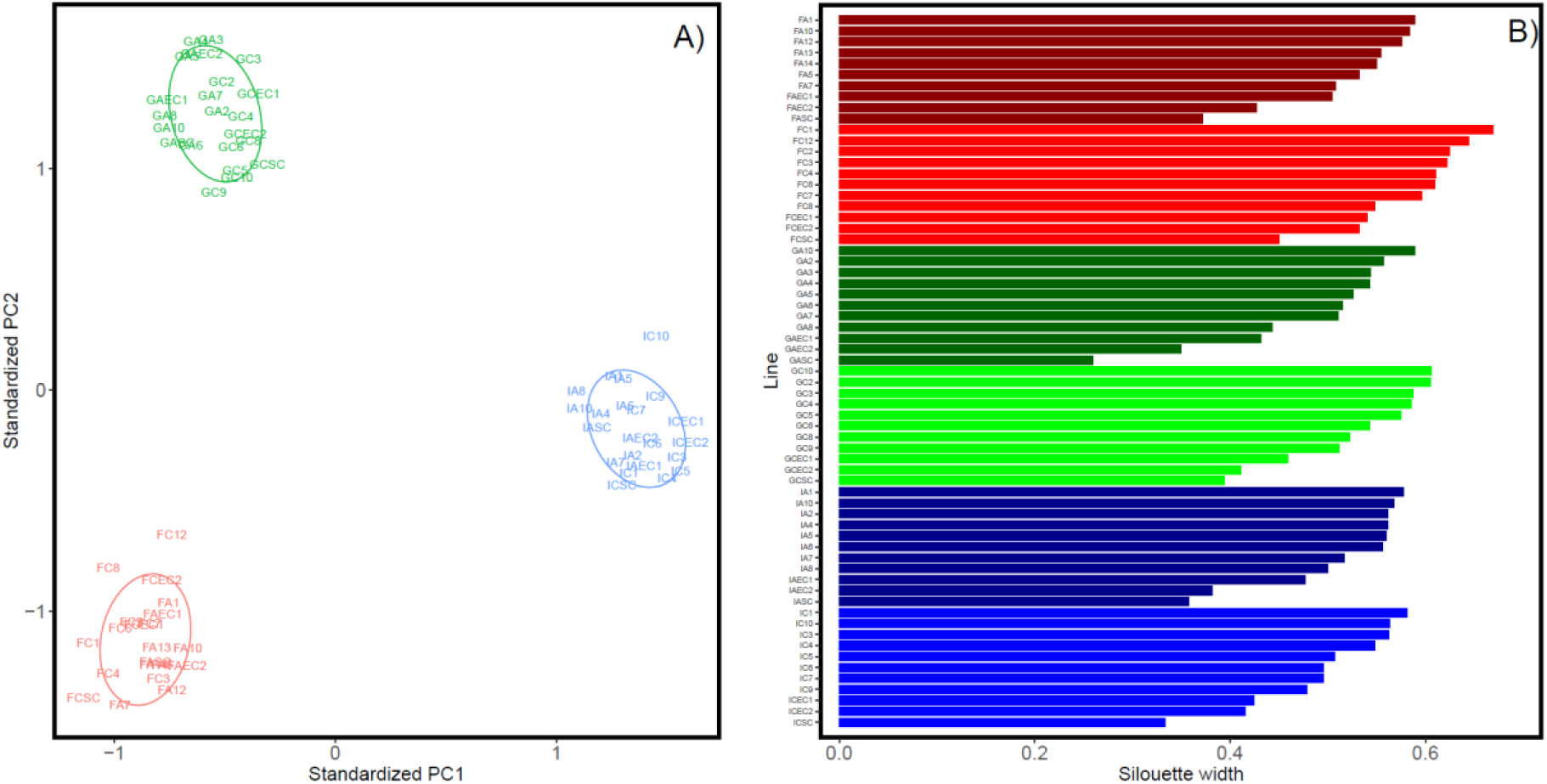
Population structure using the 100 kmers with non-zero copy number across all 65 lines. (A) Each line is plotted based on the first and second principle components axis. Lines from Finland, Germany and Israel are coloured red, green and blue, respectively. (B) k-medoids analysis using the first 10 PCs of the principal components analysis. Each cluster only contained one starting genotype (SC) and all of its descendant MA and EC lines. Dark red, red, dark green, green, dark blue and blue represents lines from genotypes FA, FC, GA, GC, IA, IC, respectively.

Our PCA results are conservative because we only examine kmers shared across all lines (including the kmers unique to each population would only increase the degree of clustering observed). We observed 92, 91 and 127 kmers, respectively, that only exist in the lines from Finland, Germany and Israel. Unsurprisingly, the average repeat unit length of these population-specific kmers were 13.8, 14.1 and 12.8 for Finland, Germany, and Israel, respectively, which are larger than the average of the 100 shared kmers. The vast majority of population-specific kmers are low in abundance, with an average copy number below 25. The exceptions are AATAGC and ACTCCT with average copy numbers of 130 in IA and 87 in IC, respectively (but which are each still present but rare in the other genotype from that region).

We found a range of 104 to 148 kmers that appeared at least twice in all SC, EC and MA lines of a particular genotype and observed 283 unique kmers across all genotypes. There were 13 highly abundant kmers with an average copy number ≥ 100 across the SC lines of all genotypes (Table 1) which ranged in length from 1 to 6 bp. In contrast, *D. pulex* has 39 repeats with an average copy number ≥ 100 (Flynn et al., 2017)and these kmers ranged in length from 1 to 20 bp. Of the highly abundant *D. magna* kmers, 12 out of 13 exist in the *D. pulex* dataset, while only 19 of the 39 highly abundant *D. pulex* kmers exist in our *D. magna* dataset. In both species, the most abundant kmer was the 1-mer A and followed by the 5-mer AACCT, but the copy number was much higher in *D. pulex* for both (Table 1). As noted previously, AACCT is likely an ancestral telomeric repeat in Arthropods that is present in several crustaceans (*D. pulicaria, Gammarus pulex* and *Penaeus semisulcatus*) and insects (Okazaki et al., 1993; Sahara et al.; Schumpert et al., 2015). On average, 30% of the total kmer base pairs were composed of AACCT in *D. magna* and 26% in *D. pulex* (Flynn et al., 2017). Kmers C, AAC and AAG had similar copy numbers between the two species, while the remaining eight high abundance kmers in *D. magna* had lower copy numbers or were absent in *D. pulex* (Table 1).

**Table 1.** Normalized copy number of highly abundant kmers for each SC line from six genotypes of *D. magna* collected from three locations, Finland (F), Germany (G) and Israel (I).

| kmer | FA | FC | GA | GC | IA | IC | *D. pulex* * |
| --- | --- | --- | --- | --- | --- | --- | --- |
| A | 21847 | 22096 | 19529 | 22181 | 12965 | 18481 | 85440 |
| AACCT | 9767 | 8083 | 4399 | 6347 | 6799 | 11390 | 48905 |
| AAG | 7325 | 6333 | 5047 | 8817 | 5178 | 7292 | 1893 |
| C | 7246 | 4892 | 2661 | 5796 | 2495 | 2870 | 4982 |
| AG | 3900 | 2528 | 4231 | 3001 | 3179 | 4687 | 376 |
| ACTAT | 2136 | 2462 | 1569 | 1678 | 1316 | 1956 | - |
| AAAAC | 988 | 1376 | 627 | 1236 | 1296 | 676 | 8 |
| AC | 622 | 567 | 604 | 624 | 948 | 971 | 269 |
| AAC | 285 | 278 | 324 | 360 | 354 | 440 | 279 |
| AGC | 177 | 168 | 209 | 180 | 167 | 150 | 55 |
| AAT | 190 | 150 | 188 | 169 | 134 | 142 | 4 |
| AT | 151 | 159 | 113 | 165 | 149 | 134 | 2 |
| AACAGG | 53 | 105 | 171 | 110 | 128 | 232 | 31 |

\*Copy number for *D. pulex* represent the mean across their six non-MA lines

For the kmers with at least two copies across all lines of a genotype, the distribution of repeat unit lengths was similar across the six genotypes (Table 2). Kmers with short lengths tend to have higher copy number than longer kmers. We observed an abundance of kmers with lengths divisible by three (i.e. 3-, 6-, 9-, 12-, 15-and 18-mers) and an abundance of 5-mers. Kmers with lengths 5, 6, 12 and 15 were also very common in *D. pulex*. However, *D. pulex* contained an abundance of 10-and 20-mers (15 and 50, respectively), which we did not observe in *D. magna*.

**Table 2.** Count and average copy number for kmers of different lengths (k) found in each genotype of *D. magna* assayed in this experiment (also with *D. pulex* data from Flynn et al. (2017)).

| *D. magna* |  |  |  |  |  |  |  | *D. pulex* |  |
| --- | --- | --- | --- | --- | --- | --- | --- | --- | --- |
| k | # kmers |  |  |  |  |  | Mean copy number | # kmers | Mean copy number |
|  | FA | FC | GA | GC | IA | IC |  | All lines |  |
| 1 | 2 | 2 | 2 | 2 | 2 | 2 | 13578 | 2 | 42954 |
| 2 | 3 | 3 | 3 | 3 | 3 | 3 | 1361 | 2 | 326 |
| 3 | 9 | 9 | 8 | 9 | 9 | 9 | 668 | 5 | 2159 |
| 4 | 8 | 6 | 3 | 3 | 5 | 6 | 31 | 4 | 267 |
| 5 | 11 | 13 | 9 | 10 | 12 | 10 | 938 | 10 | 8953 |
| 6 | 14 | 15 | 8 | 18 | 17 | 19 | 31 | 10 | 148 |
| 7 | 3 | 1 | 1 | 3 | 5 | 5 | 45 | 1 | 189 |
| 8 | 2 | 2 | 2 | 3 | 1 | 0 | 7 | 3 | 202 |
| 9 | 3 | 3 | 6 | 9 | 9 | 8 | 10 | 4 | 11 |
| 10 | 5 | 4 | 2 | 3 | 5 | 5 | 9 | 15 | 875 |
| 11 | 6 | 3 | 3 | 2 | 2 | 3 | 21 | 1 | 10 |
| 12 | 25 | 21 | 17 | 22 | 24 | 19 | 6 | 19 | 128 |
| 13 | 4 | 4 | 2 | 3 | 8 | 7 | 12 | 5 | 156 |
| 14 | 2 | 2 | 1 | 1 | 3 | 2 | 6 | 3 | 16 |
| 15 | 26 | 24 | 16 | 16 | 19 | 21 | 6 | 13 | 15 |
| 16 | 1 | 1 | 1 | 1 | 0 | 0 | 6 | 1 | 7 |
| 17 | 1 | 1 | 2 | 2 | 0 | 0 | 4 | 6 | 175 |
| 18 | 16 | 18 | 16 | 16 | 18 | 16 | 6 | 6 | 17 |
| 19 | 5 | 5 | 1 | 1 | 2 | 2 | 5 | 2 | 16 |
| 20 | 2 | 2 | 1 | 1 | 2 | 2 | 5 | 50 | 439 |

\*Mean copy number was averaged across all lines for all kmers of repeat unit length k

### Microsatellite mutation rate profiles of D. magna

For each of the six genotypes, we estimated mutation rates for kmers with at least six copies in the SC line and at least two copies in each of the EC and MA lines. The number of kmers with estimated mutation rates ranged from 60 to 79 across the six genotypes, which totaled to 144 unique kmers. Across all six genotypes, 31 kmers were present in all populations, while 20, 22 and 29 kmers were unique to genotypes of Finland, Germany and Israel, respectively. We observed that the absolute value of the mutation rate, |U_i,j_| was strongly positively correlated with the initial copy number of the kmer in the SC line for each genotype (average correlation = 0.75, Figure S1). This was expected because kmers with higher representation in the genome likely represents a larger mutational target. To remove this correlation, we divided the mutation rate of each kmer by its initial abundance to obtain an estimate of the per copy mutation rate, u_i,j_ (average correlation = 0.0031, Figure S1). It is important to note, the program we used (kseek) estimates the copy number across all arrays of a particular kmer and thus our mutation rates is an estimate of the net change in copy number due to increases and decreases at all arrays, rather than an estimate of array length changes (see Methods for details). Thus, a positive (negative) value for the genome wide or per copy mutation rate does not mean that the particular kmer only experienced increases or expansions (decreases or contractions) in copy number, rather it means that the net effect of mutation was to increase (decrease) copy number.

We observed high levels of variation in microsatellite mutation rates for *D. magna*, ranging from negative (net decrease in copy number for a given kmer) to positive (net increase in copy number for a given kmer), even among lines from the same genotypes. Across all MA lines and kmers, the genome-wide mutation rate (U_i,j_) ranged between −1103 to 2370 copies per generation and the per copy mutation rate (u_i,j_) ranged between −0.19 to 0.33 copies per initial copy number per generation. We found that mutation rates varied considerably across the six genotypes, but not consistently between genotypes of the same population. Figure 2 shows the kmer mutation rates (both U_j_, u_j_) averaged across all kmers for each genotype. For both U_j_ and u_j_, FA, FC and IC had negative mutation rates (meaning a decrease in copy number), while GA and IA had positive mutation rates, on average. GC had negative U_j_, but positive u_j_, on average, which indicates that there was one or more kmers that possessed low negative per copy mutation rates (u_j_), but were abundant enough to cause the average of U_j_ to be negative (which weights u_j_ by the abundance of kmer j).

**Figure 2.**
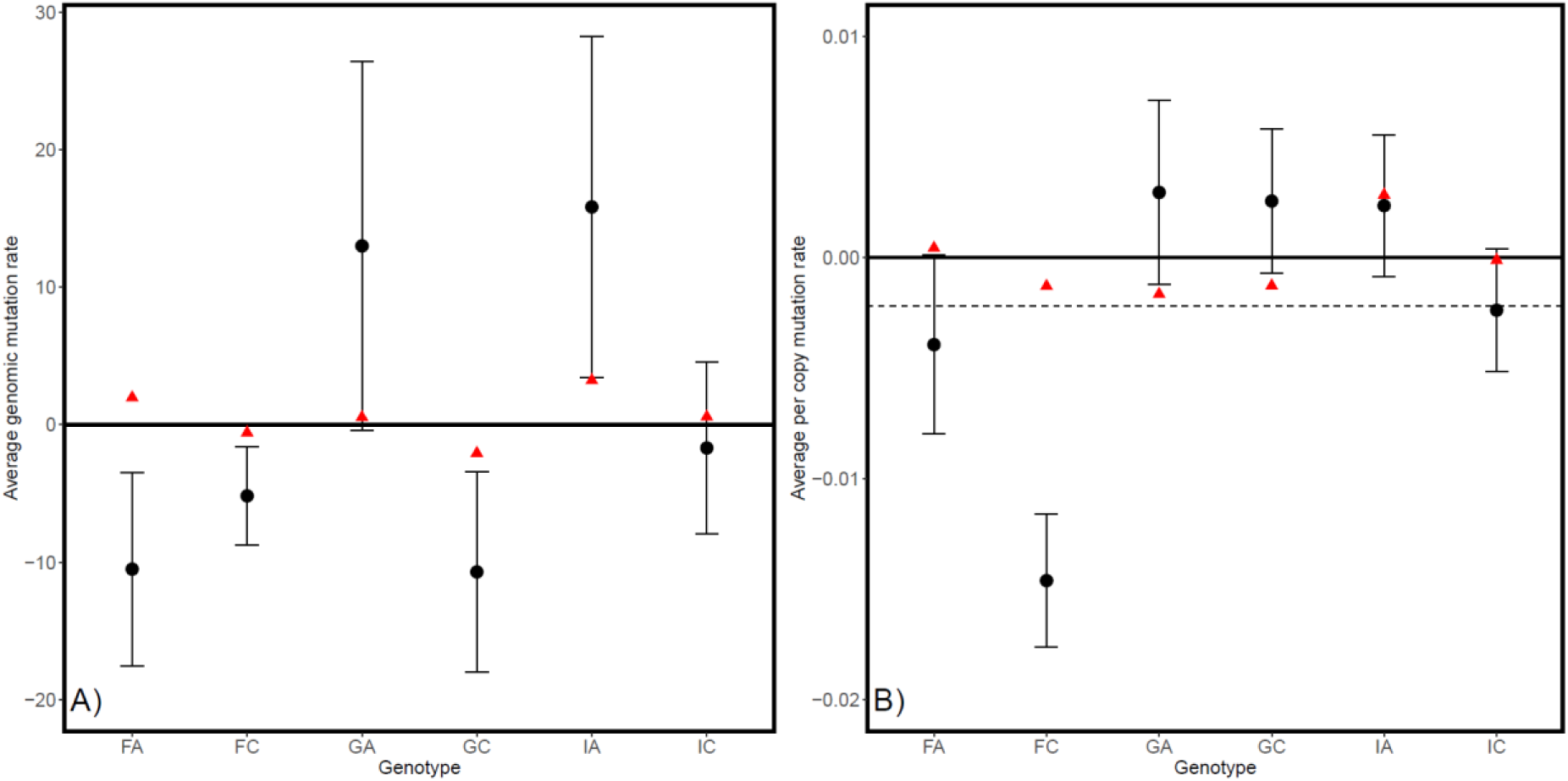
Mean (+/- SE) genomic mutation rate (A) and per copy mutation rate (B) for each genotype from six genotypes of *D. magna* collected from three locations, Finland (F), Germany (G) and Israel (I). Black circles and red triangles represent MA and EC lines, respectively. The dashed line represents the mutation rate of MA lines averaged across all genotypes.

We used the absolute value of the per copy mutation rates (|u_i,j_|) to examine the magnitude of kmer copy number change. Across all MA lines and kmers, the average absolute per copy mutation rate ranged from 0.0000042 to 0.33 and had a mean of 0.029. EC lines had a lower rate of kmer copy number change with an average 0.0049, suggesting that selection indeed constrained the rate of kmer copy number change in these large population controls.

Overall, kmer mutations rates were higher in magnitude and more variable in *D. magna* than in *D. pulex*. To compare to the *D. pulex* dataset, we applied a similar filter (requiring that each kmer considered have at least two copies in all MA lines and at least six copies across all non-MA lines (as a proxy for the ancestor)). For the 121 kmers for which we were able to estimate absolute per copy mutation rates, |u_i,j_|, the mean was 0.004 copies per copy per generation (ranging from 0 to 0.053), which is an order of magnitude lower than the average rate for *D. magna* MA lines. Only 21 of the 121 *D. pulex* kmers were present in *D. magna* (Figure S3), and the average |u_i,j_| for these 21 kmers in *D. pulex* and *D. magna* was 0.0041 and 0.031 copies per copy per generation, respectively. Furthermore, the coefficients of variation in |u_i,j_| for these 21 kmers were consistently lower in *D. pulex* than in the *D. magn*a genotypes (Figure S3).

### Mutation rate variation based on features of the kmer

Per copy mutation rates of individual kmers, u_j_, varied greatly between kmers of different lengths (Figure 3, Figure S4). Figure 3 shows the average value of u_j_ across kmers of the same length (k) for each genotype, which we define as u_j_(k). This value [u_j_(k)] can be positive or negative at most kmer lengths, depending on the particular genotype. Per copy mutation rates were most positive in 1-mers and tend to be more negative in kmers with k ≥ 8. However, fitting a linear model (lm (u_j_ ∼ k)), for each genotype did not show that per copy mutation rate was significantly correlated with kmer length (Table S1, Figure S4), likely because of the considerable variation in mutation rates even among kmers of the same length. Indeed, Kruskal-Wallis tests across kmers of the same length show significant variation in u_j_ within genotypes for most kmer lengths (Table S2, Figure S4), in addition to the variation in mutation rates at each kmer length observed between genotypes illustrated in Figure 3.

**Figure 3.**
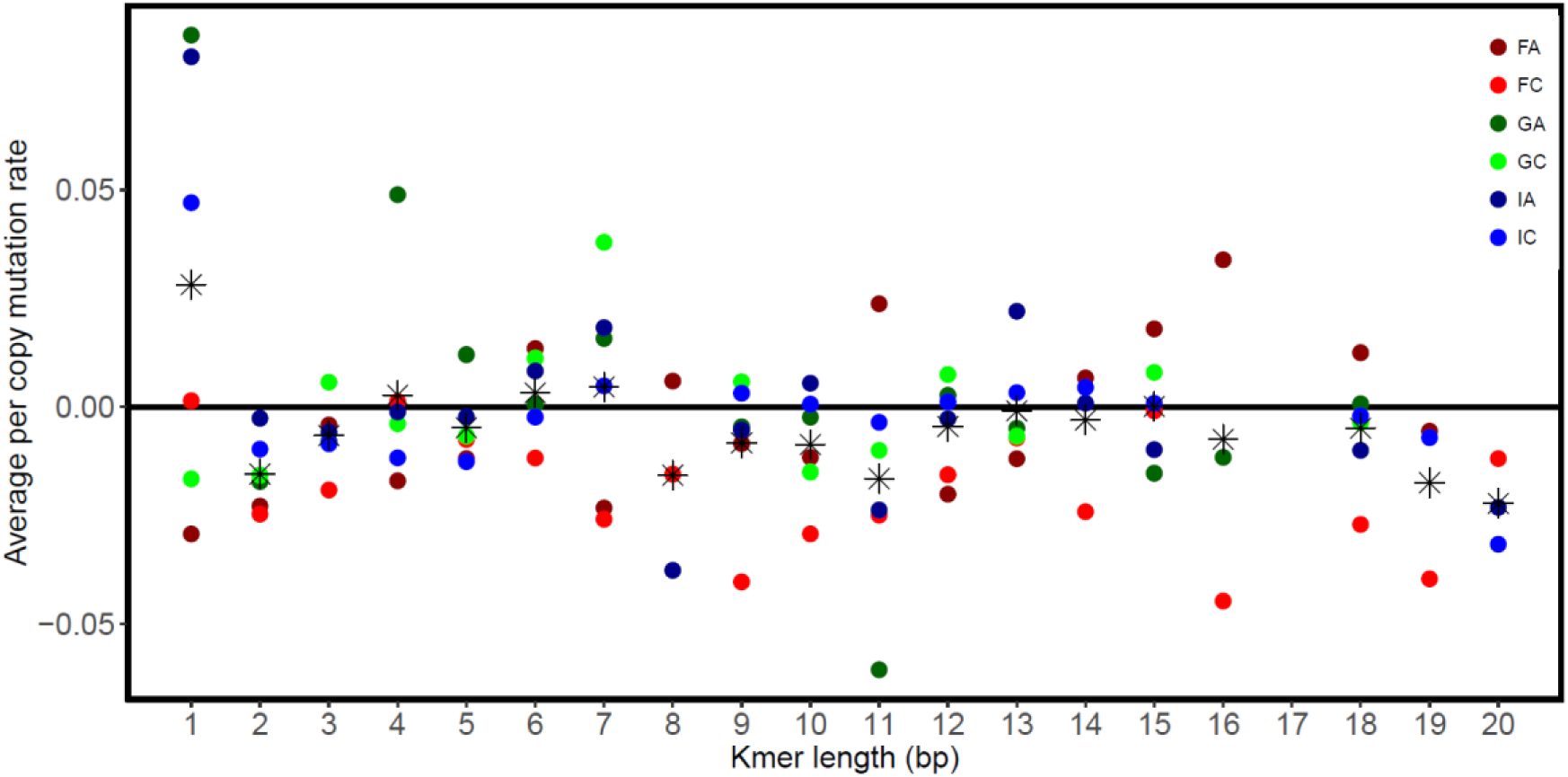
Means of kmer per copy mutation rate for each genotype and length of kmer from six genotypes of *D. magna* collected from three locations, Finland (F), Germany (G) and Israel (I). The asterisk symbol represents the average value across the six genotypes.

We also examined whether GC-content may have affected mutation rates. Kmers containing higher levels of GC may have a lower propensity to undergo mutation because GC pairs forms a more stable bond than AT pairs. To measure the propensity for mutation, we calculated the absolute values of per copy mutation rates (|u_j_|), which combines the rates of kmer copy increases and decreases, for kmers with repeat unit lengths longer than three base pairs. For each genotype, we examined the GC-content of the ten kmers with the highest and the ten with the lowest absolute per copy rates (|u_j_|; Figure 4, Table S5). We excluded kmers less than three base pairs long because these kmers will have extreme values of GC-content; including the 1-mers and 2-mers did qualitatively change our results. We observed that average |u_j_| for the kmers with the highest rates were at least four times higher than kmers with the lowest rates (Table S4). As predicted, the kmers with lower |u_j_| tend to possess higher GC-content across all genotypes. A two-way ANOVA for GC-content with genotype and mutation rate category (high vs low) as factors revealed that GC-content significantly differed between kmers with the highest and lowest mutation rates (Table S6).

**Figure 4.**
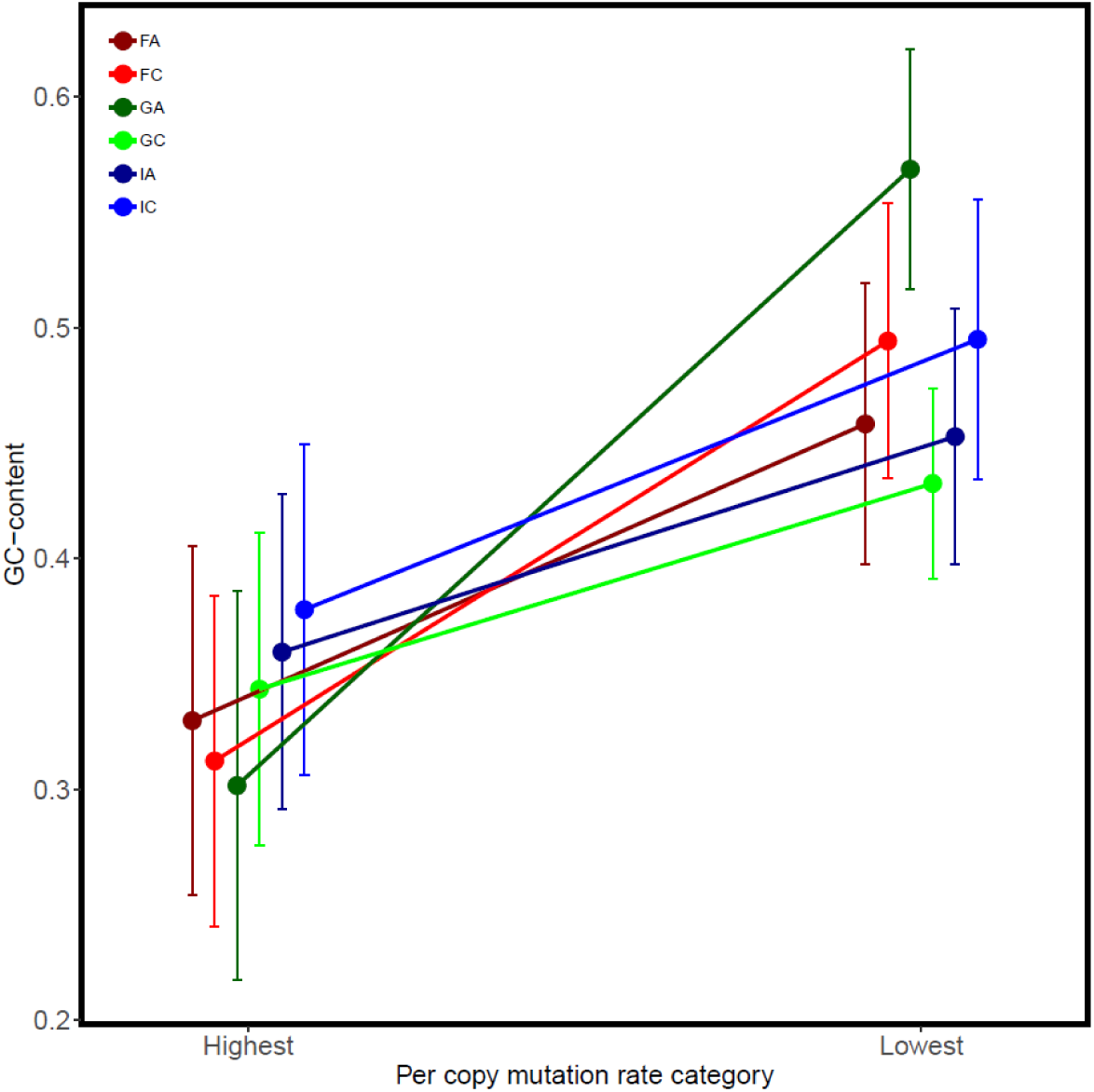
GC-content (mean +/- SE) of kmers with the top highest and lowest absolute per copy mutation rates, |uj|, from six genotypes of *D. magna* collected from three locations, Finland (F), Germany (G) and Israel (I).

### Linking variation in microsatellite landscapes and microsatellite mutation rates

The total amount of change in kmer content during mutation accumulation is not consistent within or between populations (Figure 5). Both genotypes from Finland (FA, FC) experienced a reduction in kmer content per generation, on average, while one genotype from Germany and Israel (GA and IA) experienced an increase in kmer content while the other experienced a decrease (GC and IC). Since the MA lines of GA and IA experienced the greatest increases in kmer base pairs, on average, we expected these two genotypes would contain the highest amount of kmer content, overall, but the opposite is true (Figure 5). The SC lines of GA and IA contain the lowest kmer content initially, but exhibit the highest rates of increase due to mutation. In contrast, the SC lines of FA, FC, GC and IA contained the highest kmer content and showed the greatest declines in kmer content during mutation accumulation. We tested if the change in GC-content of the microsatellite portion of the genome also varied based on starting GC-content level, but observed no significant relationship (Figure S5). Thus, instead of exhibiting strong differences based on population of origin (Figure S7) or features of individual kmers, the major differences in mutation rate profiles appear to depend on features of the genome-wide kmer content, overall.

**Figure 5.**
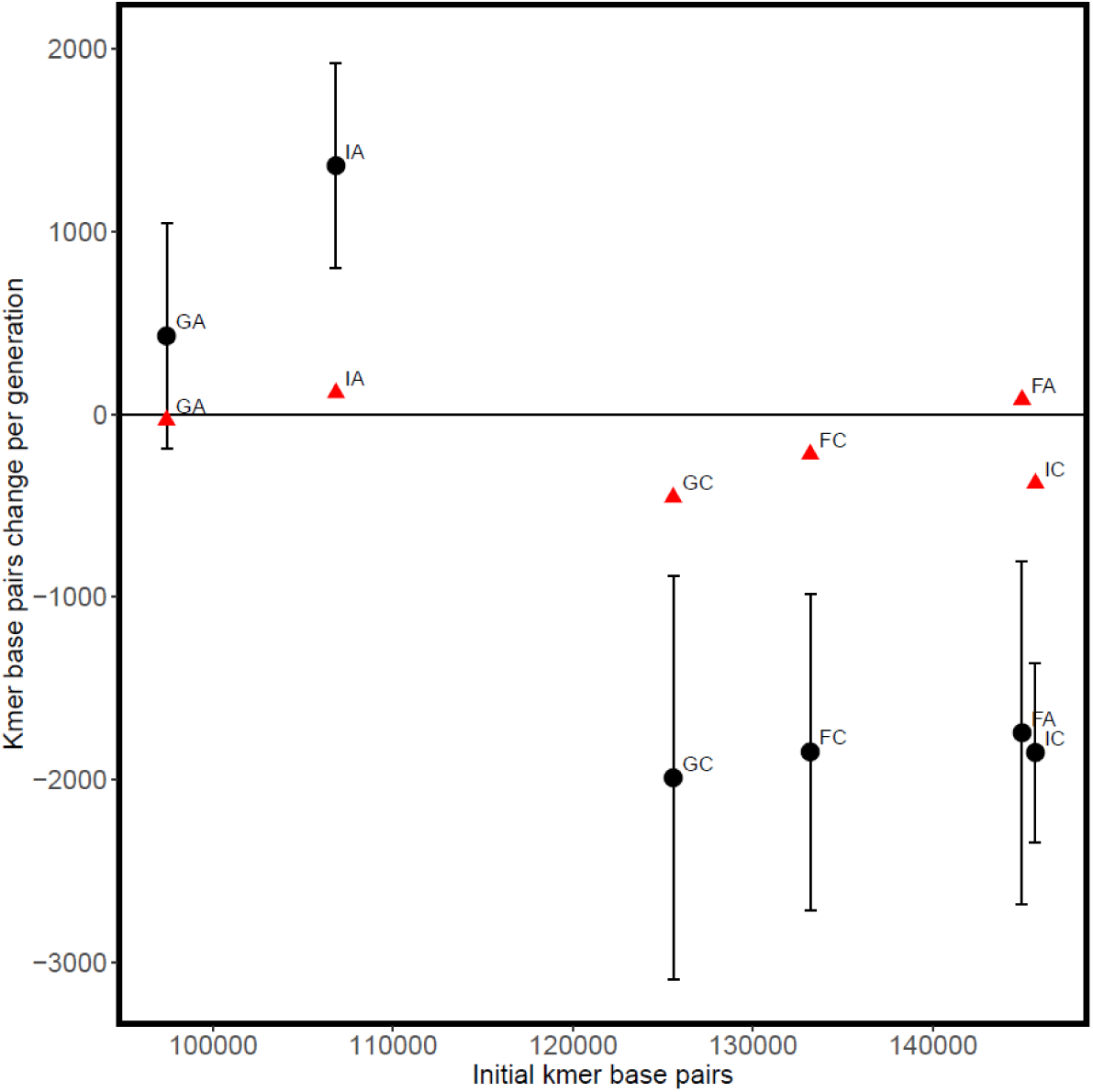
Mean (+/- SE) kmer base pair change per generation for six genotypes of *D. magna* collected from three locations, Finland (F), Germany (G) and Israel (I). Black circles and red triangles represent MA and EC lines, respectively.

As alluded to previously, average kmer mutation rates ranged from positive to negative and varied between genotypes without being consistent within populations (Figure 2). We can examine this in more detail by focusing on the 31 kmers with mutation rate estimates across all six genotypes. Kruskal-Wallis tests show significant variation in u_j_ across genotypes for all but two of the kmers (Figure 6, Table S3). If kmer mutation profiles were similar within populations, we would expect high correlations in u_j_ for genotypes from the same population, however this was not observed (Table S6). IA and IC possessed a relatively strong positive correlation (0.67) in u_j_, but they also shared a strong correlation with GA (IA-GA: 0.60, IC-GA: 0.63). Furthermore, this correlation was mainly driven by their shared high positive u_j_ for the kmer C (Figure 6). Removing the kmer C reduced the pairwise correlations (IA-IC: 0.33, IA-GA: 0.47, IC-GA: 0.40). We also performed a principal components analysis (PCA) using the u_j_ of the 31 shared kmers and did not find evidence of clustering by population-of-origin based on principle components one and two (Figure S6). Including all 144 kmers (i.e., genotype-specific kmers) in the PCA improved clustering slightly (Figure S7).

**Figure 6.**
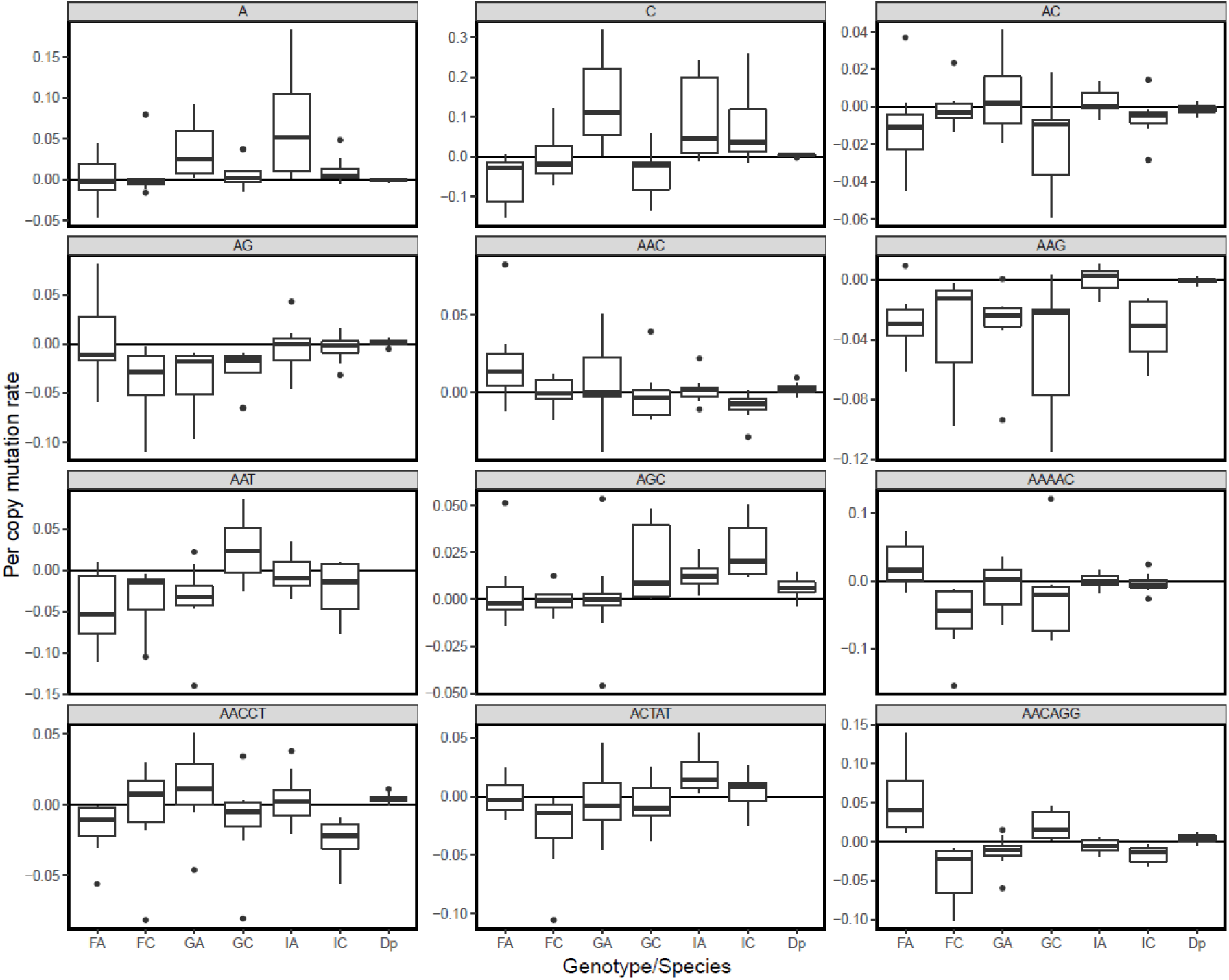
Per copy mutation rate for the 12 kmers with the highest copy number across for six genotypes of *D. magna* collected from three locations, Finland (F), Germany (G) and Israel (I). Dp represents the per copy mutation rate for *D. pulex*. Points indicate lines with per copy mutation rates less than Q1-1.5*IQR or larger than Q3+1.5*IQR. Q1, Q3 and IQR represents the first quartile, third quartile and the interquartile range, respectively. represents lines from genotypes FA, FC, GA, GC, IA, IC, respectively.

## Discussion

Repetitive regions of the genome, once overlooked, are now known to be a large and dynamic component of the genome, often responsible for large proportions of the genetic variation among individuals and species. Microsatellite loci, in particular, are known to exhibit elevated mutation rates compared to unique sequences, and have been shown to be important components of the genome in a variety of functional contexts ranging from disease risk to speciation (Gemayel et al., 2012; Hannan, 2018; Shah et al., 2010). The goal of this study was to quantify intraspecific and interspecific variation in the microsatellite landscape and microsatellite mutational dynamics in *Daphnia.* To do this, we analyzed kmers with unit lengths of 1-20 bp residing within arrays spanning at least 50 bp for six *D. magna* starting control lines (SC), 47 mutation accumulation lines (MA) and 12 non-MA existing control lines (EC). We were able to characterize the kmer profile of our *D. magna* lines based on 283 kmers, and were able to estimate mutation rates for 144 kmers. Our analysis differs from the many previous studies that examine microsatellite mutation rates because we use total kmer counts across all loci spanning at least 50 base pairs, rather than examining individual microsatellite arrays independently. Using a mutation accumulation experiment and genome-wide approach, our estimates of kmer mutation rates provide a lower-bound estimate of the net change in kmer copy number due to mutations of all types across all microsatellite loci.

### Microsatellite landscapes are distinct among genotypes, populations, and congeners

Our results clearly show distinctive microsatellite landscapes among the 6 genotypes from the three different populations of *D. magna* samples (Figure 1A and B). Using the abundance of 100 kmers presents in all lines, we observed clustering of genotypes by their population of origin (Figure 1A). In addition, within each population, we observed that MA and EC lines formed distinct clusters based on the genotype (Figure 2). These results are conservative and including kmers unique to genotypes would only strengthen the clustering. For the kmers with presence/absence polymorphism across genotypes, the vast majority were low in copy number suggesting that they arose relatively recently. In contrast, microsatellite analysis for 6 genotypes of *Chlamydomonas reinhardtii* found many kmers with hundreds of copies in some genotypes but absent or rare in others (Flynn et al., 2018). Our analyses reveal that the kmer content of *D. magna* is highly dynamic and can cause high levels of intraspecific variation, even within populations.

The microsatellite profile of *D. magna* is distinct from that of the only previously examined congener, *D. pulex* (Flynn et al., 2017), which has a much higher proportion of microsatellite content in its genome (Tables 1 and 2). *In D. pulex*, the most abundant kmers occuring in the genome tend to be shorter repeat units, with the exception of some longer repeats, such as the known arthropod telomeric sequence (AACCT)_n_ (Okazaki et al., 1993). However, there are many kmers that are unique to each species and, for kmers that are shared, many differ greatly in copy number (Table 1, 2). *D. magna* is enriched for kmers with unit lengths that are multiples of three (Table 2). It is possible that these kmer lengths are more tolerated by selection because they are less likely to cause frameshift mutations within coding regions (Metzgar et al., 2000). *D. pulex*, on the other hand, is enriched for kmers with unit lengths that are multiples of five, as has also been reported in *Drosophila melanogaster* (Wei et al., 2014). The distinctive microsatellite landscapes observed both within and between these species invites the question—do microsatellite mutation dynamics vary widely and thus explain the accumulated differences observed among genotypes, populations and species over long time periods?

### Microsatellite mutation rates vary among genotypes and between species

Mutation rates (both genome-wide increases and decreases in copy number across kmers [U_ij_] and per copy adjusted rates [u_ij_]) vary widely among genotypes (Figure 2). For per copy adjusted rates, the two genotypes collected from Finland exhibit declines in average copy number with mutation accumulation, while those from Germany exhibit increases, and the two genotypes collected from Israel split, with one genotype having an overall positive per copy mutation rate and one having a negative rate. This level of intraspecific variation in rates has not been reported previously, although this could be an artifact of most studies being conducted on only one or a few genotypes based on the assumption that mutation rate estimates can be generalized across closely-related species. One of the major take-home messages of this study is that intraspecific variation in microsatellite mutation rates is substantial, with some genotypes experiencing increases in kmer copy number and others exhibiting a decrease in kmer copies, overall. Across all kmers and genotypes, on average, copy number change was more often negative than positive (Figure 3) in *D. magna,* which is the opposite of the pattern observed in *D. pulex* (Figure S8).

### Microsatellite mutation rates as a function of kmer length and kmer content

We examined the variation in kmer abundance based on features of the kmers themselves—both length and GC-content. There was no relationship between length and copy number change (Figure 3), with one major exception–1mers exhibit the highest positive mutation rate, on average. We observed that kmers with high GC-content tend to have lower mutation rates (Figure 4 and Supplemental Table S4). This is not a surprise in that the three hydrogen bonds holding GC pairs together might be less prone to mutation than regions that are AT-rich, given there are only two hydrogen bonds between As and Ts (Calabrese and Durrett, 2003; Fan and Chu, 2007).

### The relationship between microsatellite landscape and mutation rates

We explored the relationship between initial kmer content in the genome and the mutation profiles for each genotype to determine if the microsatellite landscape could be explained by the patterns of mutation accumulation observed in the laboratory. Given that we observed a strong positive correlation between microsatellite abundance and absolute mutation rates (leading to the calculation of per copy mutation rates for our subsequent analyses), we were surprised to find that genotypes with high initial genome-wide kmer content exhibit greater decreases in microsatellite content (total change in bp) as a result of mutation than genotypes with low initial kmer content, which exhibit greater increases in the bp contributed by microsatellites during mutation accumulation (Figure 5). The context-dependency of microsatellite mutation dynamics have been reported previously, for example previous studies have shown that longer arrays tend to decrease in length whereas those with shorter arrays tend to increase (Lai and Sun, 2003; Xu et al., 2000). However, the dependency of a mutation bias towards increasing or decreasing microsatellite content on the initial total amount of microsatellites DNA has not yet been reported to our knowledge.

If starting microsatellite content does, indeed, determine the direction of copy number change as observed here *within* a species, we would predict that the extremely high microsatellite content (10-fold higher than in *D. magna*) reported for *D. pulex* in Flynn et al. (2017) would correspond with declines in copy number during mutation accumulation. This is, in fact, the opposite of what was reported–*D. pulex*, overall, shows a bias towards copy number increases (Flynn et al., 2017), while *D. magna* shows an overall bias towards decreasing copy number (illustrated by the asterisks in Figure 3; Figure S8). This observation, combined with the observation that *D. magna* exhibit a ten-fold higher overall rate of microsatellite mutation presents a genomic puzzle. It is possible that the copy number increase bias, combined with lower mutation rates, has led to a slow but tolerable accumulation of higher kmer content in the *D. pulex* genome over time. A similar explanation has been posited for plant versus animal mitochondrial genomes, where low mutation rates and a mutation bias towards insertions may have led to the tolerable accumulation of non-coding DNA resulting in, typically, much larger organellar genomes than in animals (Lynch et al., 2006).

### Comparison between D. pulex and D. magna

As mentioned, there were a few differences that limit our ability to make a direct comparison between our results for *D. magna* and those in Flynn et al. (2017) for *D. pulex* without some caveats (i.e., coverage differences and read length differences). Since kseek only counts tandem repeats spanning at least 50 bp on a read, the shorter reads and lower coverage in Flynn et al. (2017) may make it more difficult to detect kmer copies, especially of longer kmers, in their study. However, kmer content reported in *D. pulex* was actually much higher than in *D. magna* and there was no obvious bias towards detecting shorter kmers (Table 2). In fact, Flynn et al. (2017) detected many more 10-mers and 20-mers in *D. pulex* than we found in *D. magna*. An additional difference was that Flynn et al. (2017) used the average copy number of kmers in non-MA lines as a proxy for kmer estimates for the ancestral line as a baseline for calculating rates. In our dataset, non-MA lines experienced kmer content change at a much slower rate than MA lines, suggesting that while non-MA lines serve as a relatively good proxy for ancestral lines (Figure 5, S8), this could contribute to a slight underestimate of rates. Although we would not expect this to explain the order of magnitude difference in mutation rates between *D. pulex* and *D. magna* (which we observed even when only comparing their 21 shared kmers), and the differences between species in overall kmer content, the differences in the studies likely affect the sensitivity of each analysis.

### Conclusions

We observe major differences in the microsatellite landscapes accumulated over long time periods between genotypes and populations of *D. magna*, and between this species and the previously studied congener, *D. pulex*. High levels of differentiation in repeat landscapes were also previously reported, among both populations of *Drosophila melanogaster* (Wei et al., 2014) and among species of Caenorhabditid worms (Subirana et al., 2015). Given microsatellite landscapes are shaped not only by mutational inputs, but also by selection and drift, this is not a major surprise. Our results beg the question whether mutation rate differences or differential impacts of evolutionary forces play a greater role in explaining these observed differences.

While we observe high levels of variation in the mutation rates among genotypes and kmers in *D. magna*– with some exhibiting net increases in copy number and others exhibiting net decreases in copy number– the variation does not mirror the differences seen in landscapes over long time periods. In fact, genotypes with the lowest kmer content had the highest rates of copy number increase, and vice versa. Thus, it is clear the differences in microsatellite landscapes within *D. magna* are not being driven purely by mutational inputs, but instead likely reflect the interplay of mutation, selection, and drift, potentially resulting in an equilibrium with respect to individual loci (Kruglyak et al., 1998) or overall repeat content in the genome (Petrov, 2002). Overall, genotype-and kmer-specific variation in mutation rates (Figure 6) reveals a large range in terms of mutation rates in this species, and suggests that there is abundant variation upon which natural selection could act to shape mutation rates within *D. magna*.

In addition to investigating intraspecific variation and the degree to which long-term patterns of mutation accumulation would correspond to short-term mutation rates, another goal of this study was to assess the degree to which mutation rates are consistent between closely-related species. Overall, we observe much higher (10-fold) absolute microsatellite mutation rates in *D. magna* (regardless of bias towards increasing or decreasing kmer copies), than those reported for *D. pulex* (Flynn et al., *2017). Importantly, we see a mutation bias towards decreasing copy number in D. magna* (reflected by the lower overall kmer content in this species), relative to the increase bias reported for *D. pulex,* which corresponds to the much higher level of kmer content reported for that species (Flynn et al., 2017). While it is possible that a higher effective population size (N_e_) of *D. pulex* (estimated to be approximately double that of *D. magna* by (Haag et al., 2009)) allows selection to more efficiently lower the mutation rate in this species, it seems unlikely that this difference could explain the order of magnitude difference in mutation rate observed. Alternatively, the bias towards a decrease in copy number observed in *D. magna* may make the high mutation rates more tolerable, even under a similar selective regime, assuming increasing kmer content in the genome is deleterious.

It will be interesting to investigate other aspects of the mutational profile (e.g., base substitution and indel rates) of these two species in order to determine what other major differences in mutation dynamics are observed. Future studies examining the rates of contraction and expansion across kmers, as well as the differential mutability of microsatellites within, near, or far from protein-coding regions will also yield insights into the intragenomic variability in mutation rates. Although it has been common heretofore to generalize empirical estimates of mutation rates from experiments using a single genotype from a model species, the level of intra-and interspecific variation reported here suggests caution should be taken when doing so. Once we have a more complete picture of the rates, directionality, and consequences of mutation within and among species and across the genome, we will be able to better predict adaptive potential, frequencies of genetic disease, and rates of evolution for individuals and across taxa.

## Acknowledgments

We would like to thank Dee Denver and Dana Howe and the Center for Genome Research and Bioinformatics staff for their invaluable assistance with library prep and sequencing. We would also like to gratefully acknowledge our funding sources, which include an Institutional Development Award (IDeA) from the National Institute of General Medical Sciences of the National Institutes of Health under Grant #P20GM103408 (LL), Reed College sabbatical fellowship (SS), the M.J. Murdock Charitable Trust (SS), and NSF Award MCB-1150213 (SS).

## Supplementary Tables

**Table S1.** Effect of repeat unit length (k) on mutation rate from a linear model, lm(u_j_ ∼ k) for six genotypes of *D. magna* collected from three locations, Finland (F), Germany (G) and Israel (I).

| **Genotype** | **D.f.** | **Coefficient** | **P-value** |
| --- | --- | --- | --- |
| FA | 70 | 0.00132 | 0.111 |
| FC | 75 | -0.00066 | 0.232 |
| GA | 58 | -0.00140 | 0.079 |
| GC | 64 | 0.00034 | 0.648 |
| IA | 71 | -0.00103 | 0.154 |
| IC | 77 | 0.00002 | 0.969 |

**Table S2.** P-value from Kruskal-Wallis tests for the effects of kmer repeat unit length on per copy mutation rate for each unit length and genotype for six genotypes of *D. magna* collected from three locations, Finland (F), Germany (G) and Israel (I).

| k | FA | FC | GA | GC | IA | IC |
| --- | --- | --- | --- | --- | --- | --- |
| 1 | 0.0639 | 0.3446 | 0.0587 | 0.0460 | 0.6744 | 0.0587 |
| 2 | 0.0256 | 0.0221 | 0.0297 | 0.1831 | 0.1725 | 0.7926 |
| 3 | < 0.0001 | < 0.0001 | 0.0017 | < 0.0001 | 0.0012 | < 0.0001 |
| 4 | 0.0056 | 0.0140 | 0.1791 | 0.0006 | 0.0002 | 0.0290 |
| 5 | 0.0027 | 0.0002 | 0.4714 | 0.0367 | < 0.0001 | 0.0030 |
| 6 | < 0.0001 | < 0.0001 | 0.0032 | < 0.0001 | < 0.0001 | < 0.0001 |
| 7 | 0.0018 | - | - | - | 0.0005 | 0.0005 |
| 9 | - | - | 0.0660 | 0.0159 | 0.0317 | 0.4399 |
| 10 | 0.0023 | 0.0005 | 0.0117 | 0.0147 | < 0.0001 | 0.0023 |
| 11 | 0.0254 | - | - | - | - | 0.0008 |
| 12 | 0.0336 | < 0.0001 | 0.0354 | < 0.0001 | 0.0020 | < 0.0001 |
| 13 | 0.9491 | - | - | - | 0.0823 | 0.0056 |
| 14 | - | 0.2936 | - | - | - | - |
| 15 | 0.0283 | < 0.0001 | 0.0003 | 0.3305 | 0.3570 | 0.0010 |
| 18 | 0.0649 | 0.0190 | 0.0039 | 0.0003 | 0.5784 | 0.0001 |
| 19 | - | 0.2936 | - | - | - | - |

\*Only shows Kruskal-Wallis test results when there at least two kmers with length k in a genotype.

**Table S3.**
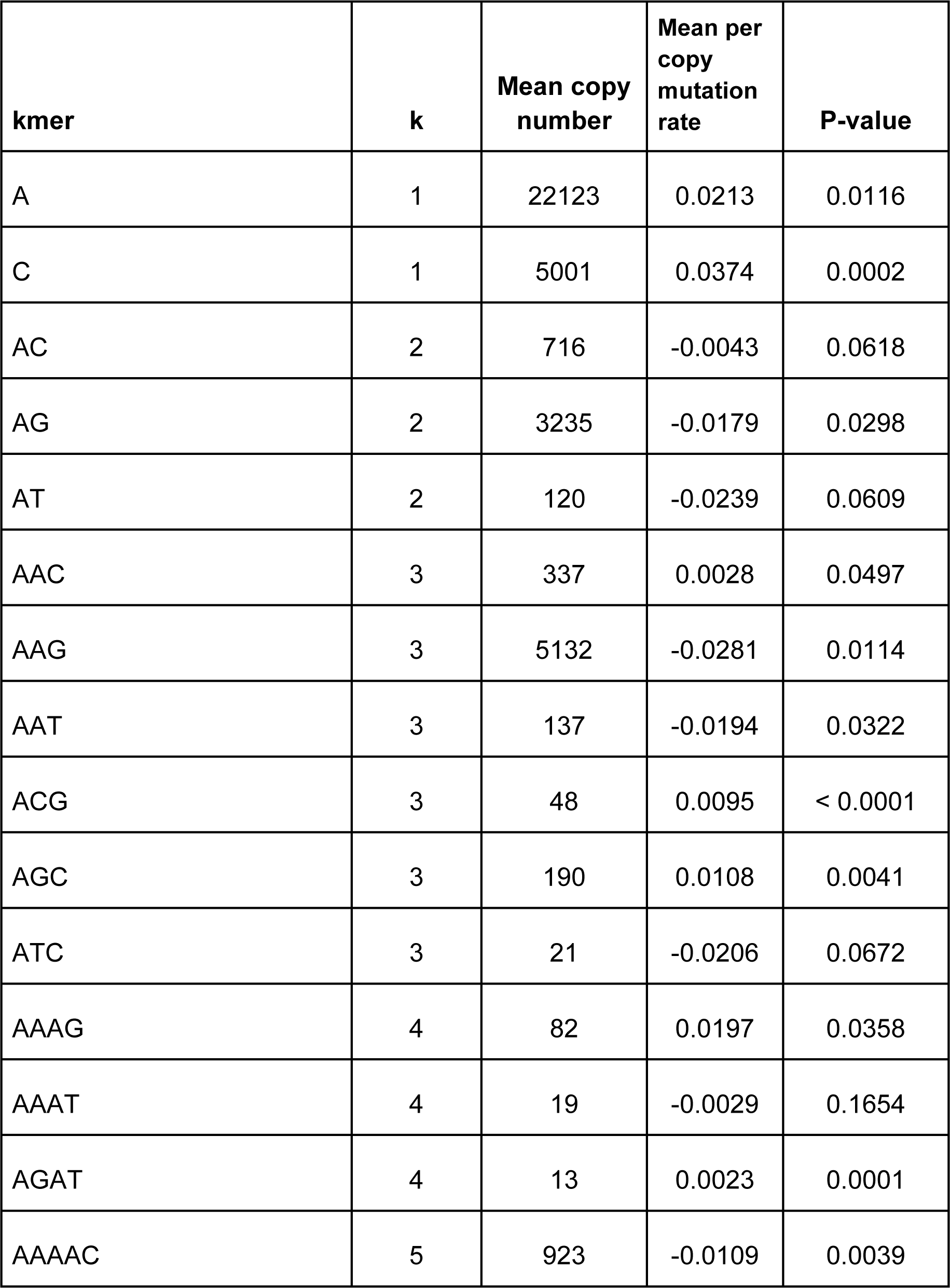
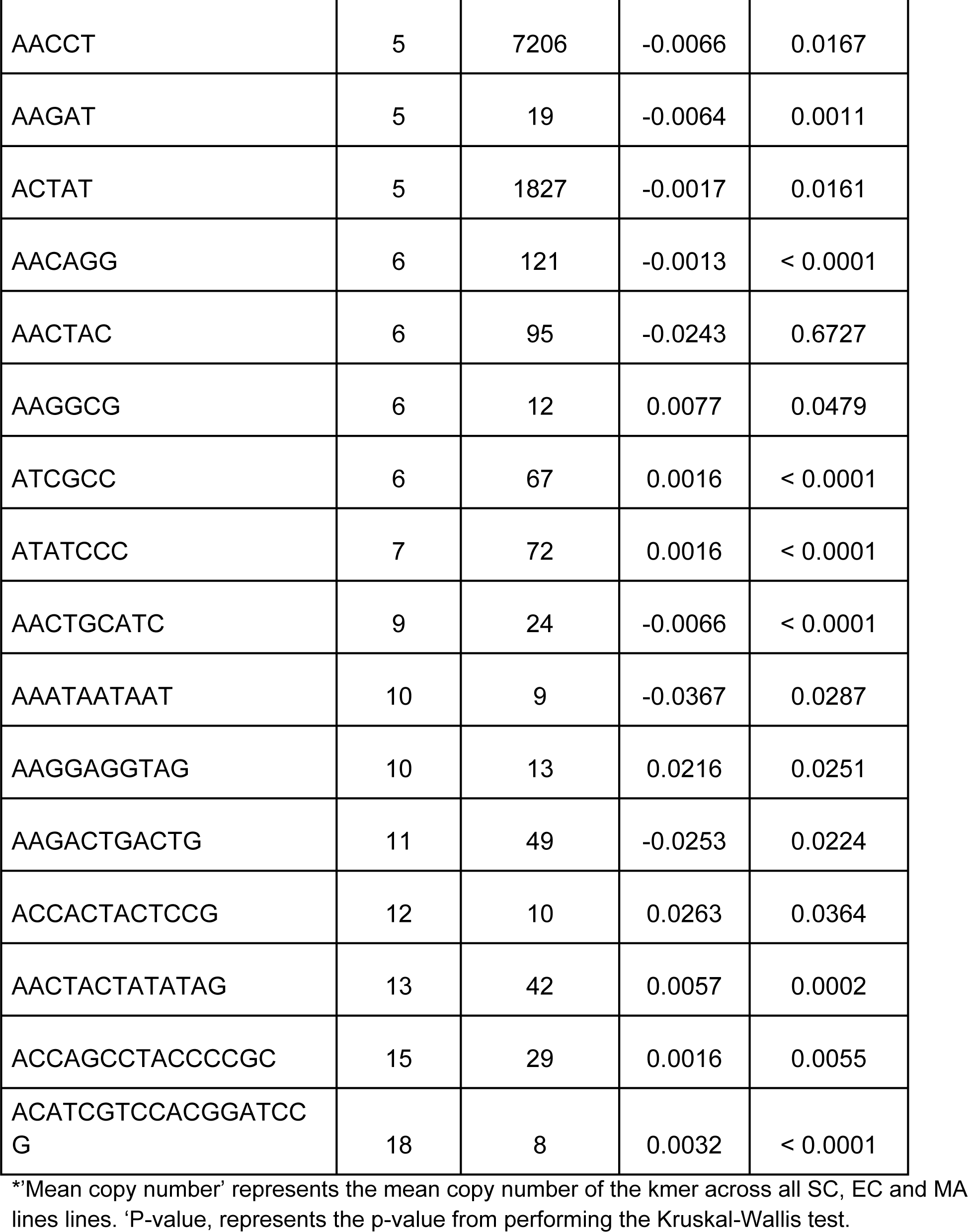
Kruskal-Wallis test of the 31 kmers with mutation rate estimates across all six genotypes of *D. magna* in this study.

**Table S4.** Statistics for the kmers with ten highest and ten lowest |u_j_| for each of six genotypes of *D. magna* collected from three locations, Finland (F), Germany (G) and Israel (I).

| Genotype | Category | Mean k | Mean GC content | Mean $\vert u_j\vert $ |
| --- | --- | --- | --- | --- |
| FA | Highest | 7.6 | 0.33 | 0.071 |
| FA | Lowest | 10.4 | 0.46 | 0.013 |
| FC | Highest | 8.1 | 0.31 | 0.056 |
| FC | Lowest | 9.4 | 0.49 | 0.012 |
| GA | Highest | 7.1 | 0.30 | 0.064 |
| GA | Lowest | 12.5 | 0.57 | 0.015 |
| GC | Highest | 6.7 | 0.34 | 0.055 |
| GC | Lowest | 8.7 | 0.43 | 0.012 |
| IA | Highest | 7.8 | 0.36 | 0.050 |
| IA | Lowest | 7.3 | 0.45 | 0.009 |
| IC | Highest | 8.0 | 0.38 | 0.044 |
| IC | Lowest | 11.4 | 0.49 | 0.010 |

\*k represents the kmer length, GC represents the proportion of base pairs that are GC in the kmer

**Table S5.** Two-way ANOVA results testing the relationship between genotype and absolute per copy mutation rate category (high vs low) and GC-content of kmers in *D. magna*.

| Factor | D.f. | SumSq | MeanSq | F-value | P-value |
| --- | --- | --- | --- | --- | --- |
| Genotype | 5 | 0.043 | 0.0085 | 0.202 | 0.9612 |
| Category | 1 | 0.641 | 0.6405 | 15.177 | 0.0002 |
| Genotype:Category | 5 | 0.115 | 0.0231 | 0.547 | 0.7405 |
| Residuals | 108 | 4.558 | 0.0422 |  |  |

\*Category high and low represents kmers with the 10 highest and 10 lowest absolute per copy mutation rate, $|u_j|$ , respectively.

**Table S6.** Pairwise correlations of per copy mutation rates among the 31 kmers shared across the six genotypes of *D. magna* collected from three locations, Finland (F), Germany (G) and Israel (I).

|  | FA | FC | GA | GC | IA | IC |
| --- | --- | --- | --- | --- | --- | --- |
| FA | 1 | 0.29 | 0.00 | 0.17 | -0.13 | 0.08 |
| FC |  | 1 | 0.45 | 0.15 | 0.41 | 0.43 |
| GA |  |  | 1 | -0.15 | 0.60 | 0.63 |
| GC |  |  |  | 1 | -0.07 | 0.01 |
| IA |  |  |  |  | 1 | 0.67 |
| IC |  |  |  |  |  | 1 |

**Table S7.** Total number of reads for each line.

| Line | Num. of reads | Line | Num. of reads | Line | Num. of reads |
| --- | --- | --- | --- | --- | --- |
| FA10 | 117276453 | GA10 | 67872281 | IA10 | 77469618 |
| FA12 | 111118502 | GA2 | 74933433 | IA1 | 76339146 |
| FA13 | 98944469 | GA3 | 78929079 | IA2 | 82762562 |
| FA14 | 118671293 | GA4 | 78740281 | IA4 | 94985444 |
| FA1 | 100759904 | GA5 | 79320460 | IA5 | 110594565 |
| FA5 | 108817148 | GA6 | 71148348 | IA6 | 100880878 |
| FA6 | 92034278 | GA7 | 76557733 | IA7 | 78383275 |
| FASC | 88590180 | GA8 | 58644700 | IA8 | 90086901 |
| FC12 | 68037364 | GASC | 72730208 | IASC | 76126994 |
| FC1 | 67658931 | GC10 | 75779178 | IC10 | 83523929 |
| FC2 | 70000714 | GC2 | 96690080 | IC1 | 84683921 |
| FC3 | 87568787 | GC3 | 74165418 | IC3 | 79042669 |
| FC4 | 72157642 | GC4 | 100073584 | IC4 | 89897718 |
| FC6 | 70810578 | GC5 | 107328336 | IC5 | 64317095 |
| FC7 | 82224489 | GC6 | 92860991 | IC6 | 67608868 |
| FC8 | 71891429 | GC8 | 106121415 | IC7 | 67236965 |
| FCSC | 93661924 | GC9 | 105658620 | IC9 | 70519081 |
| FAEC1 | 92815980 | GCSC | 71010993 | ICSC | 64756748 |
| FAEC2 | 101739760 | GAEC1 | 83641911 | IAEC1 | 77480400 |
| FCEC1 | 79655226 | GAEC2 | 70749011 | IAEC2 | 84786933 |
| FCEC2 | 92606026 | GCEC1 | 119412606 | ICEC1 | 89041032 |
|  |  | GCEC2 | 107606136 | ICEC2 | 89387923 |

## Supplementary Figures

**Figure S1.**
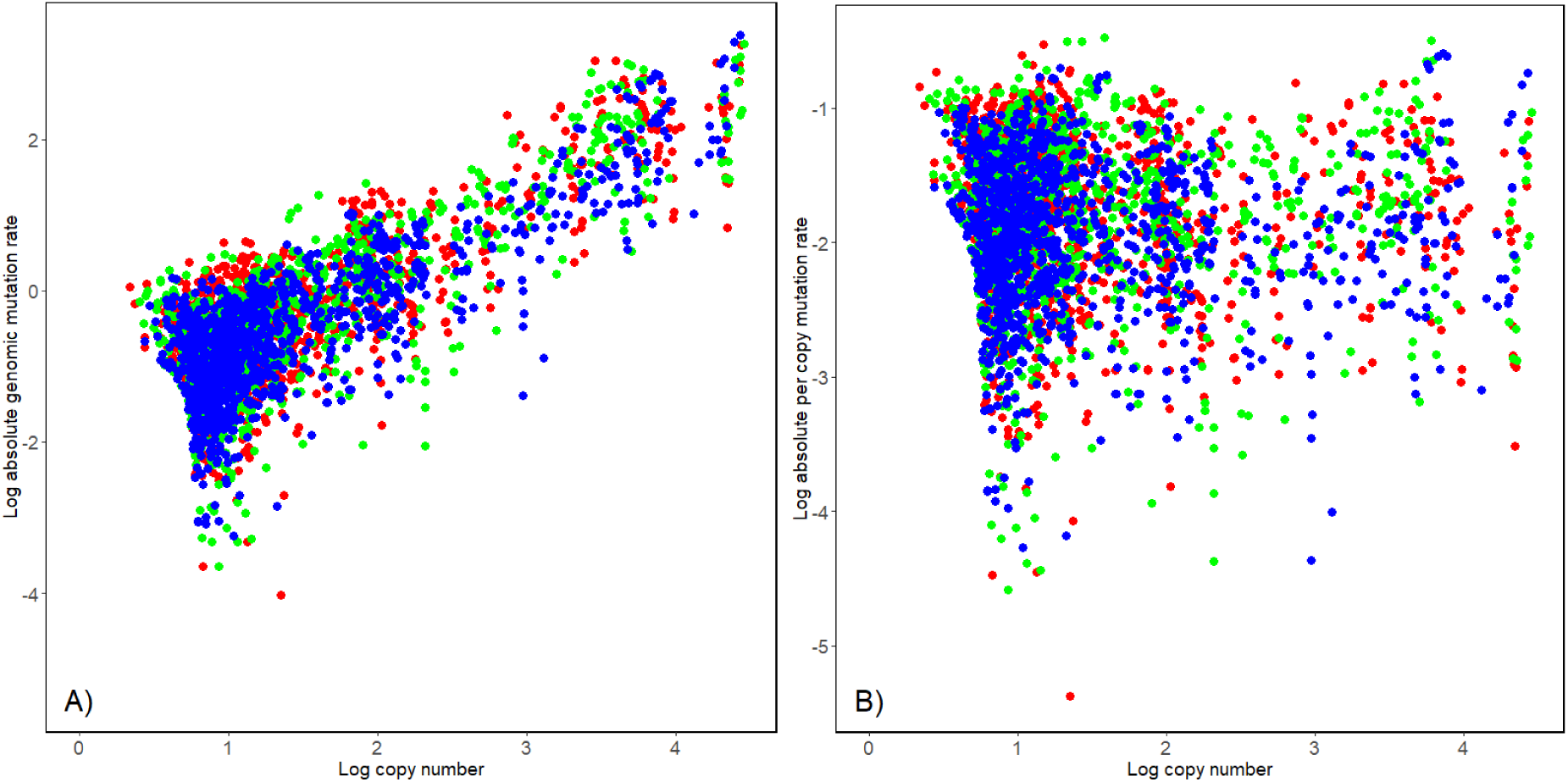
Absolute genomic mutation rate (A) and absolute per copy mutation rate (B) plotted against initial copy number for each kmer from each genotype. Red, green and blue represents genotypes from Finland, Germany and Israel, respectively.

**Figure S2.**
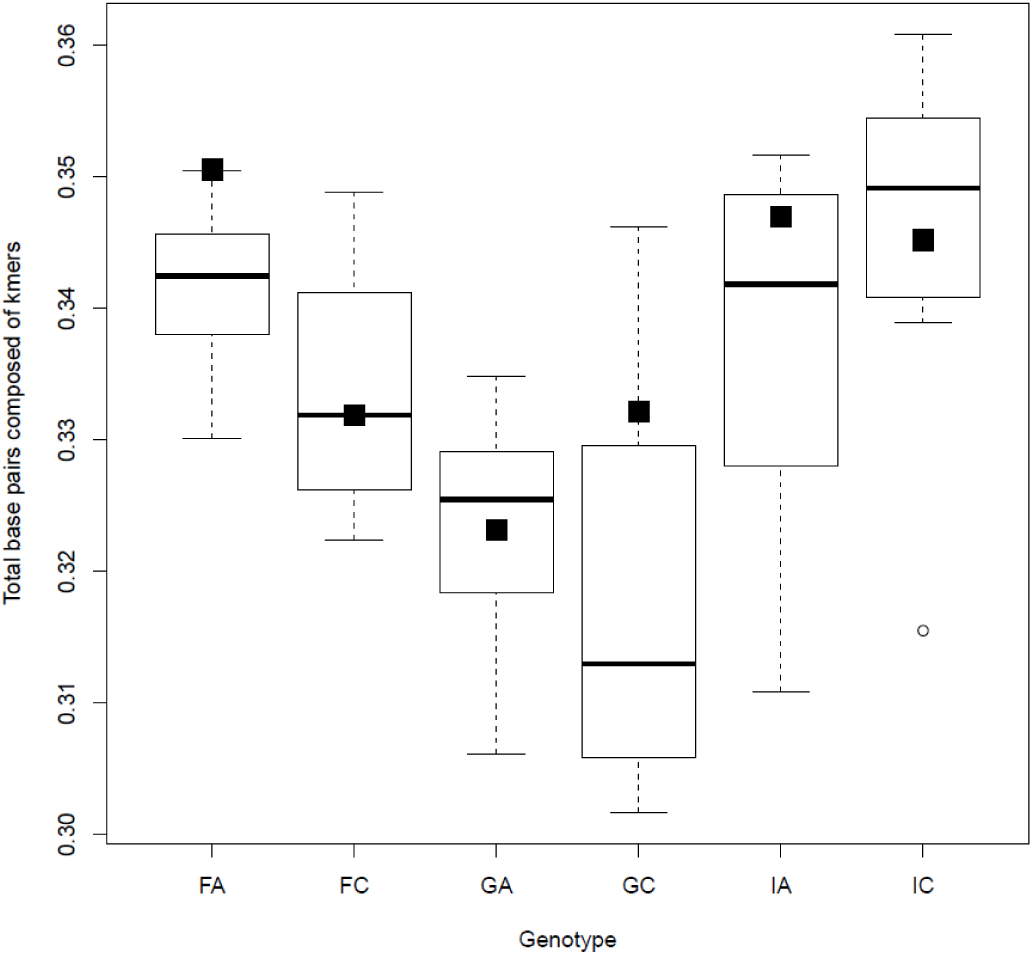
Total base pairs composed of kmers for SC and MA lines of each genotype. Total base pairs of MA lines represented by the boxplots; white circles represent lines outside 1.5 times of the interquartile range. Total base pairs for SC lines represented by the black squares. Data shown for each of six genotypes of *D. magna* collected from three locations, Finland (F), Germany (G) and Israel (I).

**Figure S3.**
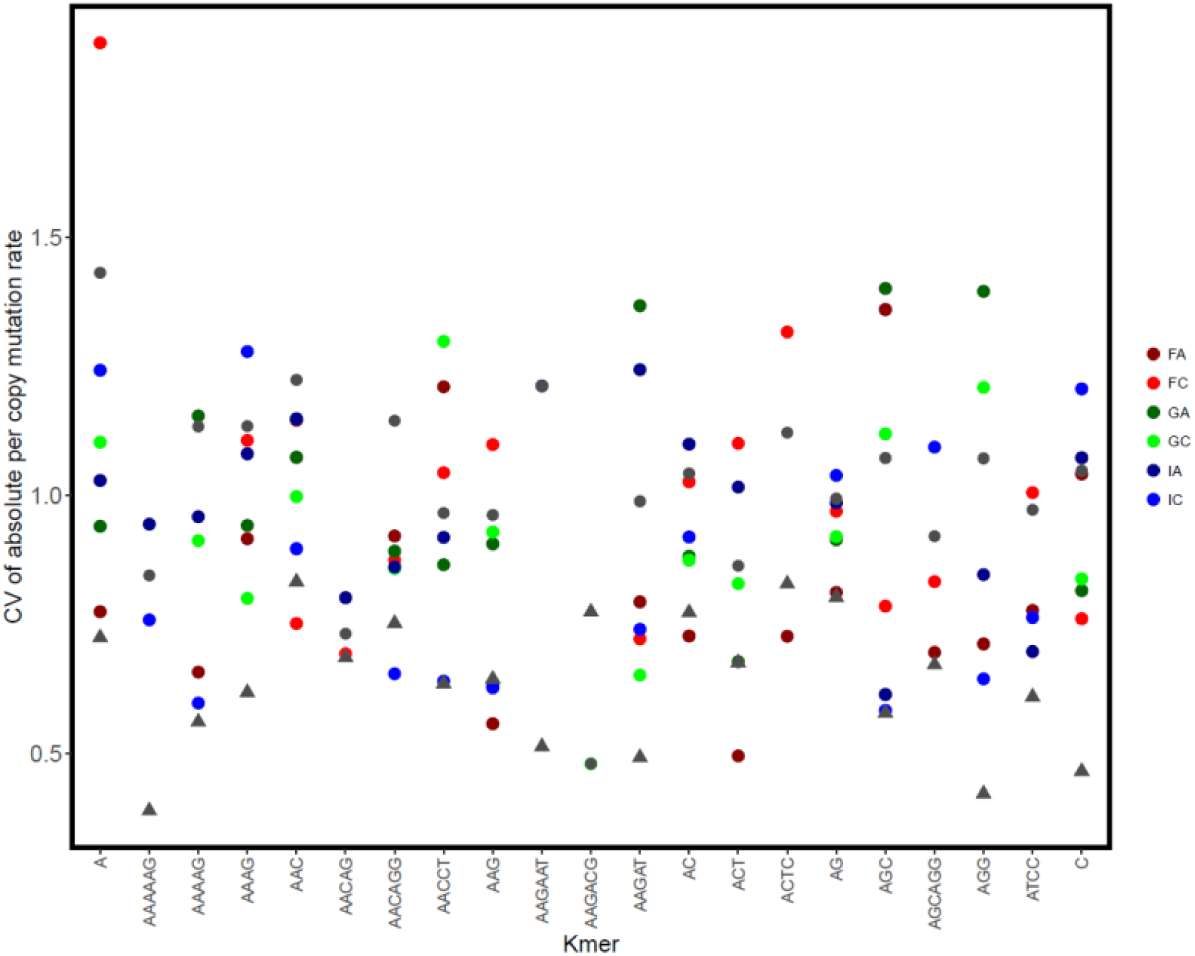
Coefficient of variation in |u_i,j_| for each *D. magn*a genotype (circle) and for *D. pulex* (grey triangle) for the 21 kmers shared across species. Grey circles represent the coefficient of variation across all six genotypes of *D. magna* collected from three locations, Finland (F), Germany (G) and Israel (I).

**Figure S4.**
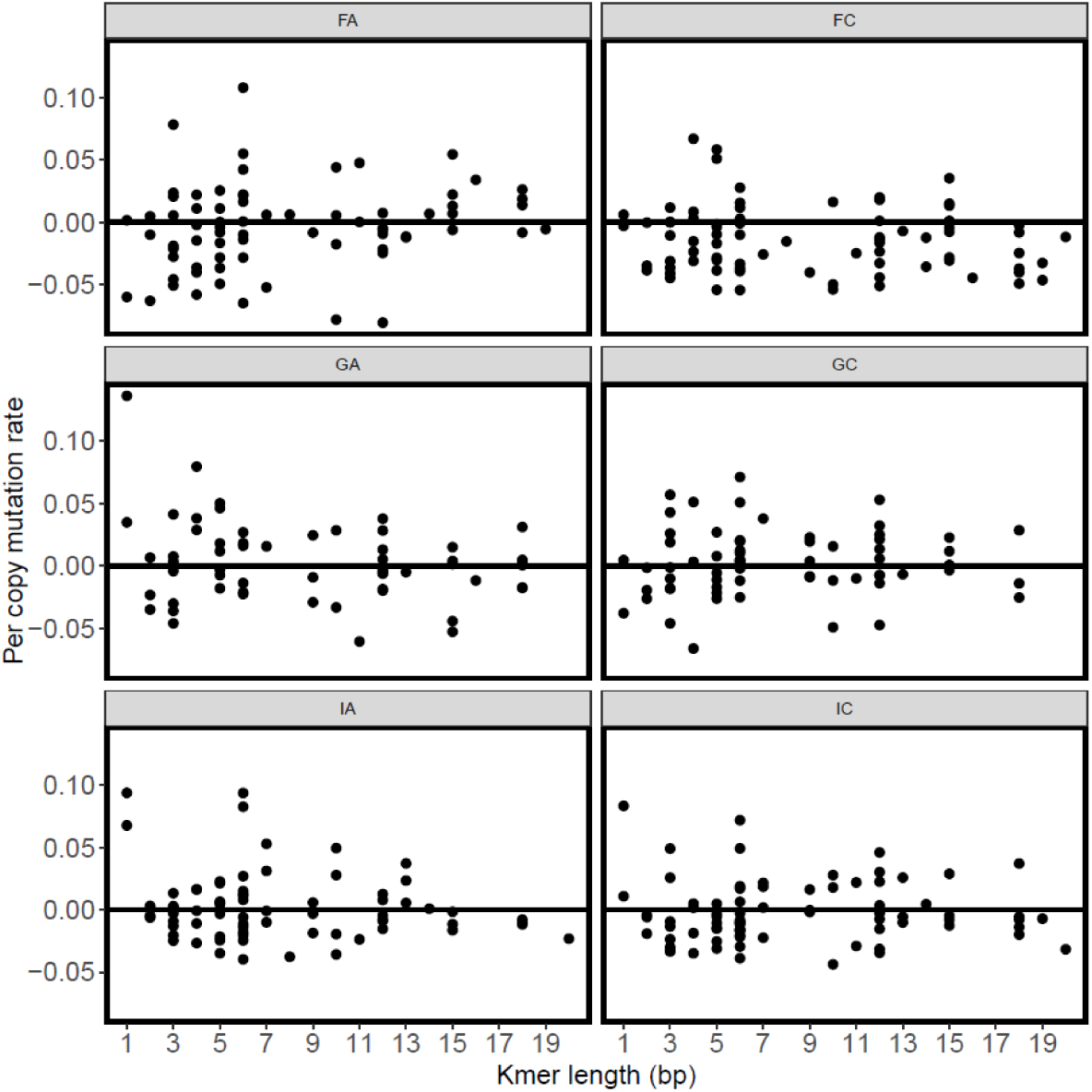
Per copy mutation rate of kmers j (u_j_) plotted against kmer lengths for six genotypes of *D. magna* collected from three locations, Finland (F), Germany (G) and Israel (I).

**Figure S5.**
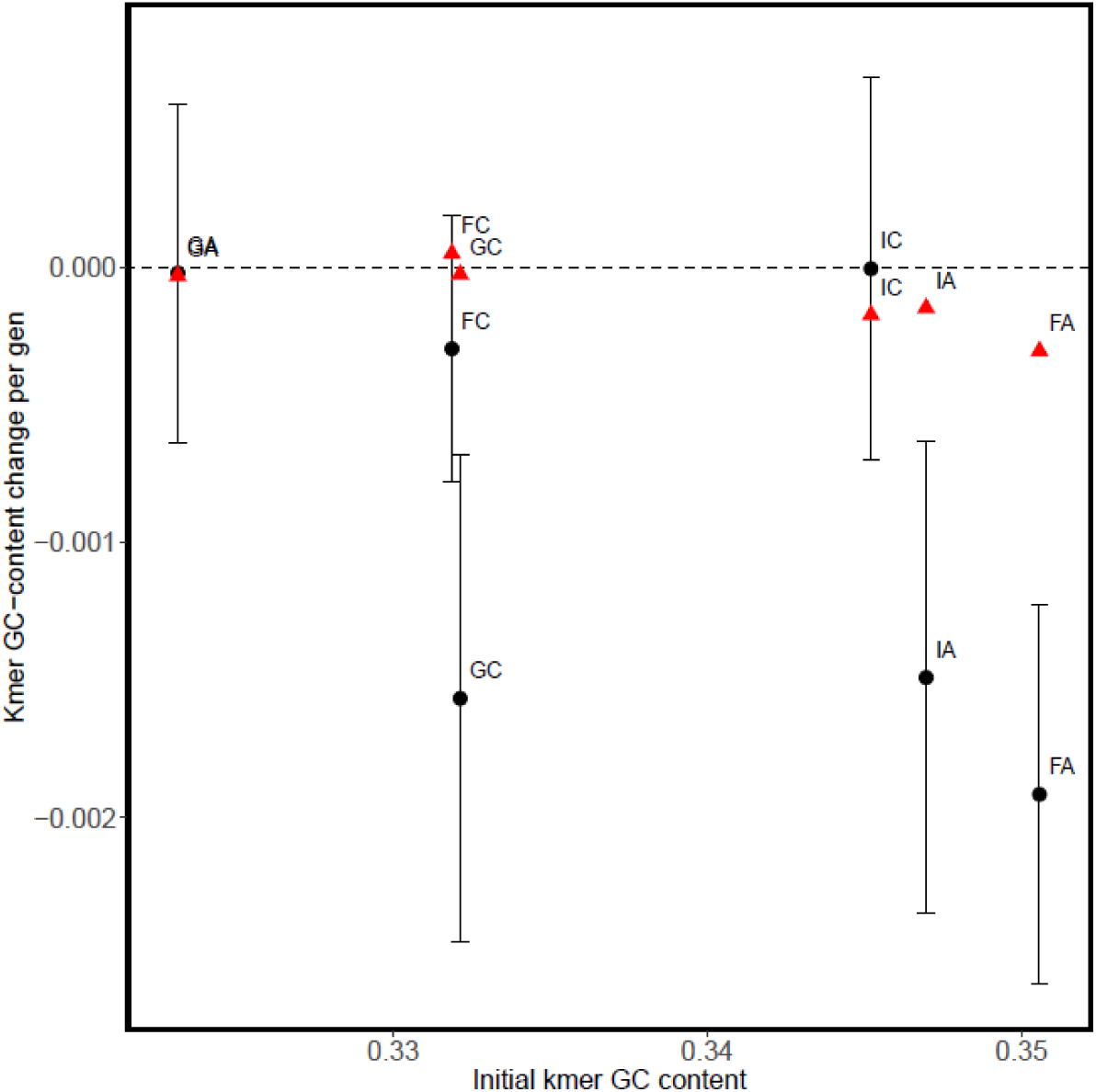
Mean (+/- SE) kmer GC-content change plotted against initial kmer GC-content for all six genotypes of *D. magna* collected from three locations, Finland (F), Germany (G) and Israel (I). Black circles and red triangles represent MA and EC lines, respectively.

**Figure S6.**
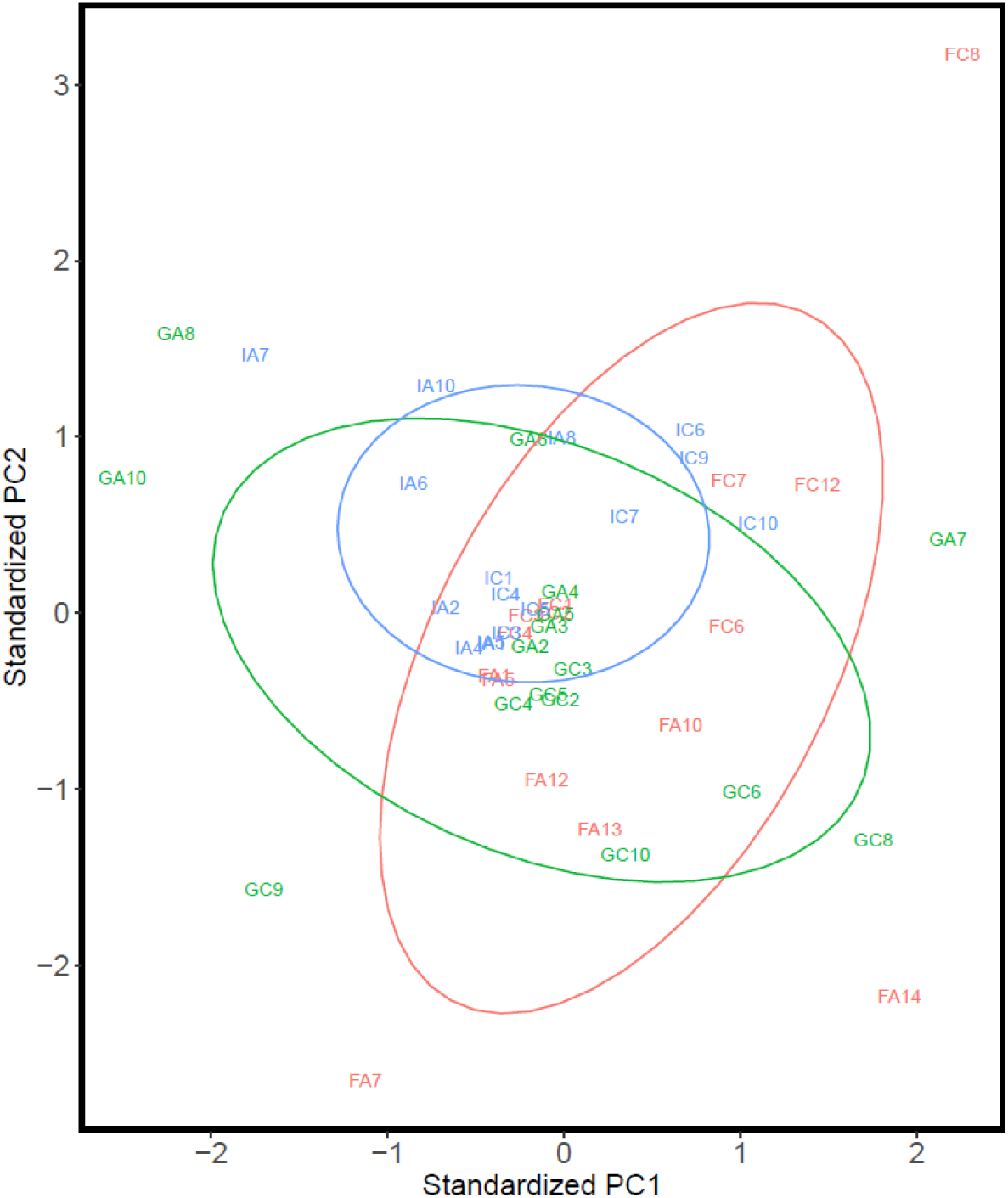
Population structure using u_j_ for the 31 kmers with mutation rate estimates for all six genotypes of *D. magna* collected from three locations, Finland (F; red), Germany (G; green) and Israel (I; blue). Each MA line is plotted based on the first and second principal components axis.

**Figure S7.**
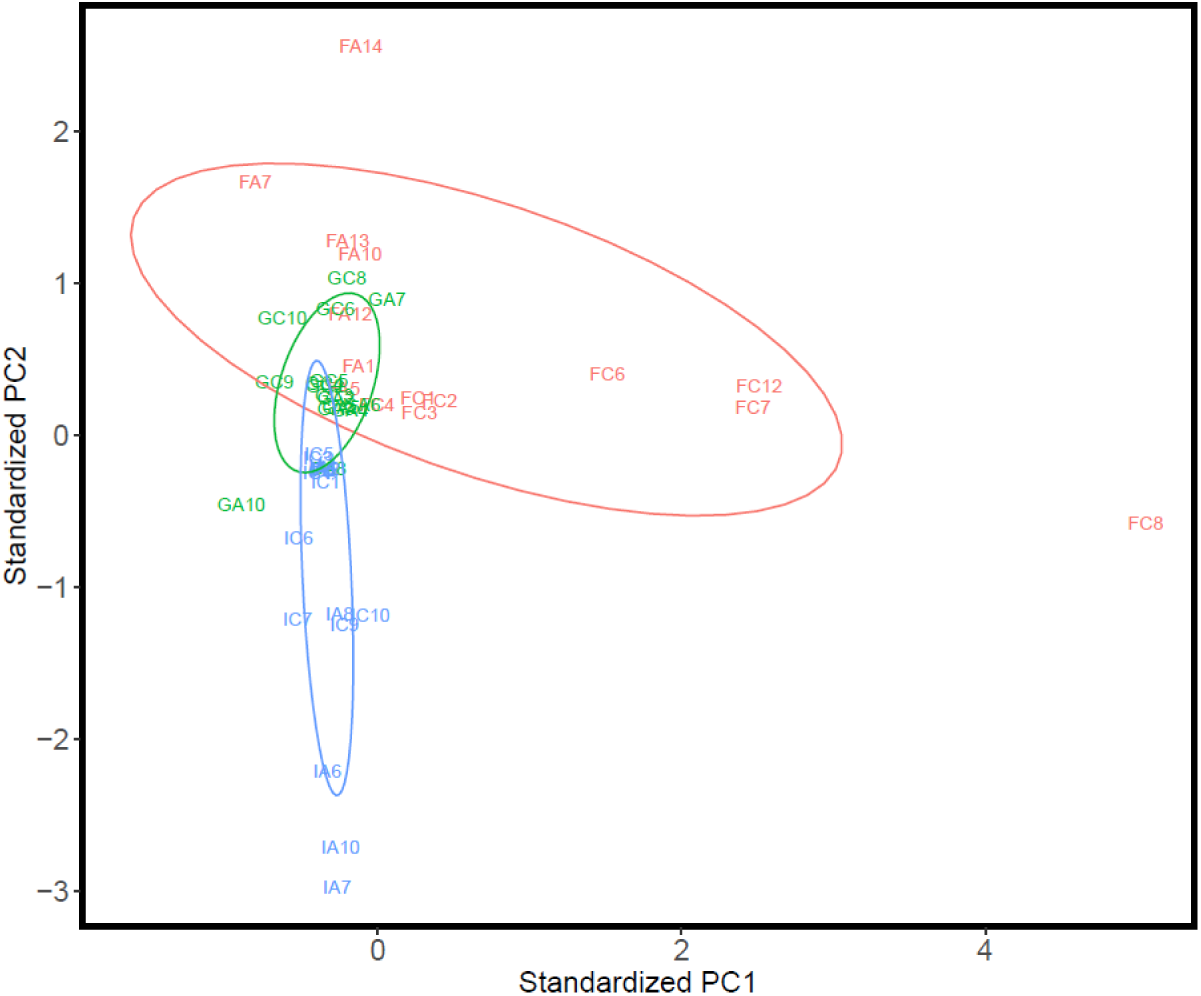
Population structure using u_j_ for all 144 kmers with mutation rate estimates for all six genotypes of *D. magna* collected from three locations, Finland (F; red), Germany (G; green) and Israel (I; blue). Each MA line is plotted based on the first and second principal components axis. If there was no mutation rate estimate for a kmer in a particular genotype, we set u_j_ as 0. Each MA line is plotted based on the first and second principle components axis.

**Figure S8.**
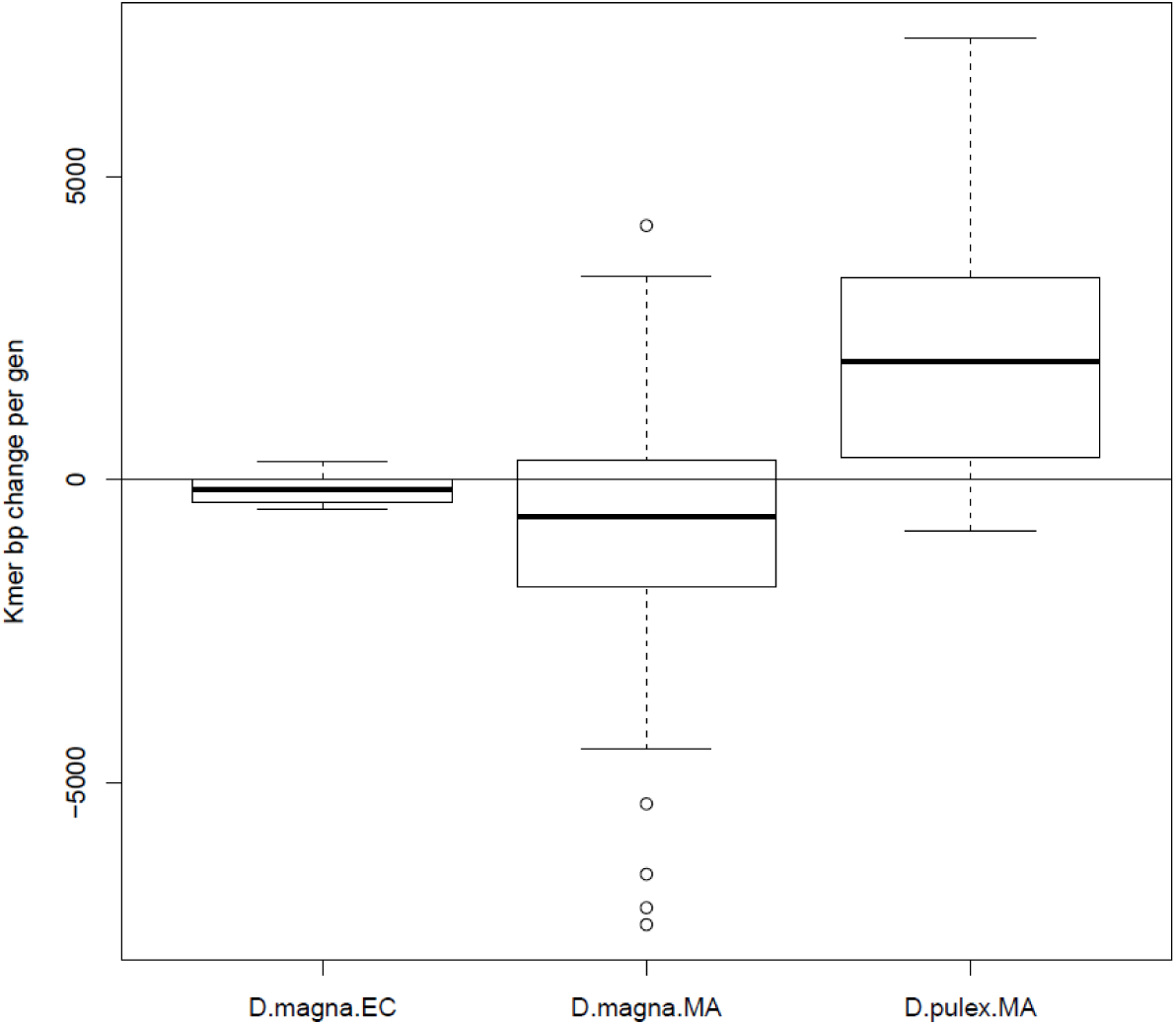
Change in total kmer content (bp) per generation for *D. magna* EC lines, *D. magna* MA lines and *D. pulex* MA lines.

